# Influence of metal ions on the isothermal self-assembly of DNA nanostructures

**DOI:** 10.1101/2024.11.04.621977

**Authors:** Arlin Rodriguez, Bharath Raj Madhanagopal, Kahini Sarkar, Zohreh Nowzari, Johnsi Mathivanan, Hannah Talbot, Vinod Morya, Ken Halvorsen, Sweta Vangaveti, J. Andrew Berglund, Arun Richard Chandrasekaran

## Abstract

DNA nanostructures are typically assembled by thermal annealing in buffers containing magnesium. We demonstrate the assembly of DNA nanostructures at constant temperatures ranging from 4 °C to 50 °C in solutions containing different metal ions. The choice of metal ions and the assembly temperature influence the isothermal assembly of several DNA motifs and designed three-dimensional DNA crystals. Molecular dynamics simulations show more fluctuations of the DNA structure in select monovalent ions (Na^+^ and K^+^) compared to divalent ions (Mg^2+^ and Ca^2+^). A key highlight is the successful assembly of DNA motifs in nickel-containing buffer at temperatures below 40 °C, otherwise unachievable at higher temperatures, or using an annealing protocol. DNA nanostructures isothermally assembled in different ions do not affect the viability of fibroblasts, myoblasts, and myotubes and or the immune response in myoblasts. The use of ions other than the typically-used magnesium holds key potential in biological and materials science applications that require minimal amounts of magnesium.

---

DNA is highly programmable for constructing nano-to-micrometer scale structures.^1^ A key advantage of DNA nanotechnology is that it offers custom-designed shapes with accurate placement of guest moieties such as biomolecules, cells, and nanoparticles, enabling biophysical,^2,3^ structural^4,5^ and biomedical^6,7^ applications. DNA nanostructure assembly typically involves a thermal annealing protocol in which DNA strands are heated in a specific buffer to a high temperature (90-95 °C) and cooled slowly at specific rates. For use as frameworks for guest molecules, a DNA nanostructure is first thermally annealed, and the temperature-sensitive guests are attached later at a lower temperature, in a two-step process.^7–9^ Assembly of DNA nanostructures under isothermal conditions is desirable for such scaffolding purposes and to reduce the need for thermal annealing instruments, allowing nanostructure preparation in low-resource settings. In prior works, 3D DNA origami structures have been assembled at a constant temperature of 55 °C,^10^ 2D structures from single-stranded tiles have been assembled at the physiological temperature of 37 °C,^11,12^ and 2D DNA origami and tile-based 2D arrays have been assembled at 25 °C.^13^ Further, additives such as formamide^14,15^ and betaine^16^ have been used to improve the assembly yields of DNA nanostructures under isothermal conditions. In addition to cargo loading advantages, assembly at constant temperatures also allows studying hybridization kinetics of DNA nanostructure assembly.^17,18^

Another important factor for DNA self-assembly is the choice of counter-ion. While most DNA nanostructure assembly strategies use Mg^2+^, the use of other ions instead, or in addition to, Mg^2+^ is useful to enhance the biostability of DNA nanostructures,^19^ increase assembly yields,^20^ and tune the plasmonic properties of DNA-nanoparticle complexes.^21^ DNA nanostructures have been assembled using thermal annealing protocols in low-magnesium buffers,^22^ by substituting Mg^2+^ with Na^+^ or by using organic cations such as spermidine^23,24^ or ethylenediamine instead of metal ions.^25^ Our own recent work showed the magnesium-free assembly of DNA nanostructures by replacing Mg^2+^ with other monovalent and divalent ions, where the structures were thermally annealed.^19^ There has not been much work on the assembly of DNA nanostructures at constant temperatures using ions other than Mg^2+^, with only one recent study demonstrating isothermal assembly in Na^+^ ions.^13^ Here, we demonstrate the assembly of DNA nanostructures at constant temperatures in solutions containing a variety of monovalent (Na^+^, Li^+^, K^+^) and divalent (Mg^2+^, Ca^2+^, Sr^2+^, or Ni^2+^) ions (**Fig. 1**). We used molecular dynamics simulations to understand the differences in the interaction of select ions with a model DNA nanostructure. We further show the biological feasibility of using DNA nanostructures assembled in these different ions by analyzing the cell viability in three different cell types, including fibroblasts, myoblasts, and myotubes, and studied the immune response of myoblasts to one of the model DNA nanostructures. To our knowledge, this is the first study on isothermal assembly and biological feasibility of DNA nanostructures assembled in a variety of counter ions.

**Fig. 1.**
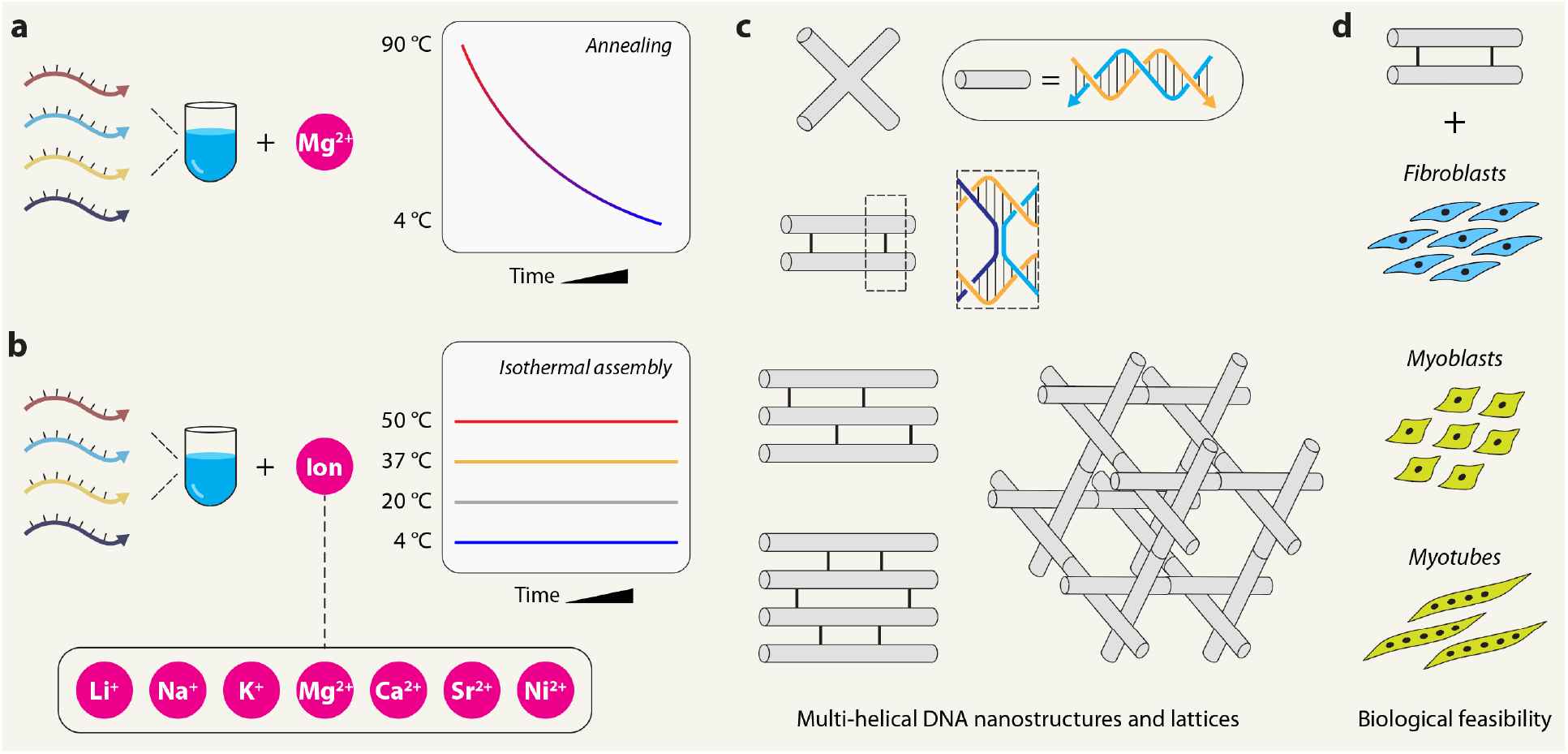
DNA nanostructure assembly. (a) Illustration of the typical annealing protocol used to assemble DNA nanostructures in magnesium-containing buffers. (b) Illustration of isothermal assembly of DNA nanostructures at constant temperatures in different cations used in this study. (c) Model nanostructures used in this study. (d) Biological feasibility analysis in different cell types.

## Results

### Isothermal assembly of double crossover DNA motif in calcium

To study isothermal assembly, we first chose Ca^2+^ as the counter ion and a DNA double crossover motif as a model nanostructure (**Fig. 2a, Supplementary Fig. 1**). The motif contains two adjacent double-helical domains connected by two antiparallel crossovers separated by an odd number of half-turns (16 bp, abbreviated DX-O).^26^ We chose this structure as a model system based on our recent work, where we assembled it in various ions using a thermal annealing protocol.^19^ We first assembled the DX-O motif in the typically used tris-acetate-EDTA (TAE) buffer containing 12.5 mM Mg^2+^ using a thermal annealing protocol and validated assembly using non-denaturing polyacrylamide gel electrophoresis (PAGE) (**Fig. 2b**). Then, we confirmed the assembly of DX-O by thermal annealing in TAE buffer containing different concentrations of Ca^2+^, instead of Mg^2+^, and observed similar assembly yields for the two metal ions (**Fig. 2c-d, Supplementary Fig. 2**). To analyze isothermal assembly, we mixed the component DNA strands (**Supplementary Table 1**) in 1× TAE containing different concentrations of Ca^2+^ (10 to 100 mM) and incubated the DNA solution at a constant temperature of 4 °C, 20 °C, 37 °C, or 50 °C for 3 hours. We analyzed the samples using non-denaturing PAGE and normalized the assembly yield to that of the sample annealed in 1× TAE containing 12.5 mM Mg^2+^ (**Fig. 2e, Supplementary Fig. 3**). While the assembly of DX-O was observed in all the temperatures tested, isothermal assembly at 37 °C and 50 °C resulted in relatively pure products, and partially folded low molecular weight structures were found in samples assembled at 4 °C and 20 °C. The isothermal assembly yields in 50 and 100 mM Ca^2+^ (at 50 °C) were comparable to samples annealed in similar Ca^2+^ ion concentrations, while samples assembled isothermally in 10 and 25 mM Ca^2+^ showed slightly lower yields compared to the annealed samples (**Fig. 2c,e**). We also observed that the assembly yields increased with time at all temperatures when assembled in 10 or 100 mM Ca^2+^, with rapid assembly at 37 °C and 50 °C (**Fig. 2f-g, Supplementary Fig. 4**). The pH of the buffer did not affect the yields at 50 °C while at other temperatures, the assembly yields were slightly higher at alkaline pH than at acidic pH (**Fig. 2h, Supplementary Fig. 5**). This observation was consistent with samples prepared using a thermal annealing protocol in TAE containing Mg^2+^ or Ca^2+^ (**Supplementary Fig. 6**).

**Fig. 2.**
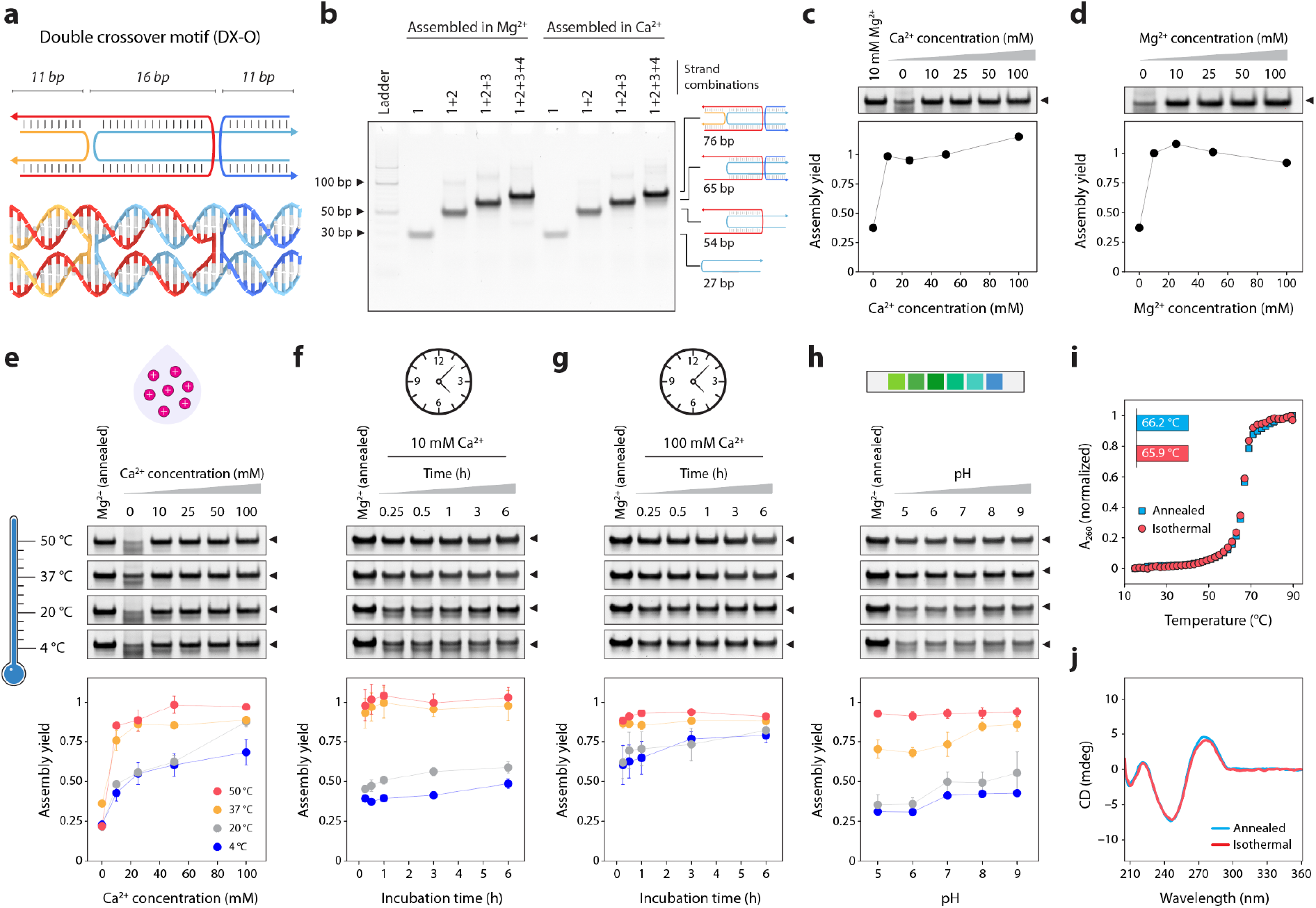
Isothermal assembly in calcium. (a) Scheme and molecular model of the DX-O motif. (b) Stepwise assembly of the DX-O motif in Mg^2+^ and Ca^2+^ using a thermal annealing protocol. (c) PAGE analysis and quantified assembly yield of DX-O assembled in different concentrations of Ca^2+^ using a thermal annealing protocol. (d) PAGE analysis and quantified assembly yield of DX-O assembled in different concentrations of Mg^2+^ using a thermal annealing protocol. (e) Isothermal assembly of DX-O at different temperatures (shown by the gradient) in buffer containing different Ca^2+^ concentrations. (f) Time series of assembly at different isothermal temperatures in 10 mM Ca^2+^. (g) Time series of assembly at different isothermal temperatures in 100 mM Ca^2+^. (h) Isothermal assembly of DX-O in different pH. (i) Thermal melting curves obtained from DX-O motif annealed in 50 mM Ca^2+^ or isothermally assembled in 50 mM Ca^2+^ at 50 °C. (j) CD profiles obtained from DX-O motif annealed in 50 mM Ca^2+^ or isothermally assembled in in 50 mM Ca^2+^ at 50 °C. Data represent mean and error propagated from standard deviations of experiments performed in triplicates.

Following the assembly optimization, we characterized the isothermally assembled DX-O motif using UV melting and circular dichroism (CD) spectroscopy. We observed that the melting temperature of DX-O assembled at 20 °C did not vary significantly with increasing Ca^2+^ concentrations (**Supplementary Fig. 7, Supplementary Table 2**). The melting temperatures of annealed and isothermally assembled DX-O (in 50 mM Ca^2+^ at 50 °C) were similar, with a Tm of 66.2 °C and 65.9 °C, respectively (**Fig. 2i**). The CD spectrum of annealed and isothermally assembled DX-O were also similar (**Fig. 2j**). These melting and CD results were consistent with earlier reports of the DX-O motif annealed in Mg^2+^,^19^ indicating proper assembly in Ca^2+^ using the isothermal procedure.

### Isothermal assembly of other DNA nanostructures in calcium

To test the isothermal assembly of other DNA nanostructures in Ca^2+^, we chose structures of increasing complexity: a 4-arm junction based on an immobile branched DNA,^27^ a double crossover DNA motif with antiparallel crossovers separated by even number of half-turns (21 bp, abbreviated DX-E),^26^ a 3-helix DNA motif that consists of three adjacent double helical domains connected by four crossovers^28^ and a 4-helix DNA motif that consists of four DNA double helical domains connected by a total of six crossovers (**Fig. 3a, Supplementary Fig. 8, Supplementary Tables 2-5**).^29^ These motifs have been used in constructing 1D^30^ and 2D arrays,^28,29^ as templates for nanowires,^31^ in algorithmic self-assembly,^32^ and for site-directed enzyme catalysis.^33^ We first confirmed proper assembly of the four structures in 1× TAE buffer containing Mg^2+^ or Ca^2+^ using a thermal annealing protocol (**Supplementary Fig. 9**), followed by isothermal assembly at 4 °C, 20 °C, 37 °C, or 50 °C in 1× TAE containing 10, 25, 50, or 100 mM Ca^2+^. For the 4-arm junction, assembly yields were similar in all the temperatures (83-95%), with slightly higher assembly yields with increasing Ca^2+^ concentration (**Fig. 3b, Supplementary Fig. 10**). For the DX-E, assembly yields were higher at 37 °C and 50 °C (**Fig. 3c, Supplementary Fig. 11**). For the more complex 3-helix and 4-helix motifs, we observed a temperature-dependent improvement in the assembly yields. Further, assembly yields increased with increasing Ca^2+^ concentrations for the 3-helix motif at 4 °C and 20 °C (18-25% in 10 mM Ca^2+^ to 50-56% in 100 mM Ca^2+^), but this effect was less pronounced at 37 °C (55% in 10 mM to 67% at 100 mM Ca^2+^) (**Fig. 3d, Supplementary Fig. 12**). Isothermal assembly at 50 °C in all Ca^2+^ concentrations was comparable to the assembly yield of the structure annealed in Mg^2+^. We observed higher assembly yields with higher isothermal temperatures for the 4-helix motif as well, with the highest assembly yields of 77-89% at 50 °C (**Fig. 3e, Supplementary Fig. 13**). The melting temperatures and CD profiles were comparable for structures assembled in 50 mM Ca^2+^ isothermally at 50 °C and using the thermal annealing protocol (**Fig. 3f** and **Supplementary Fig. 14**).

**Fig. 3.**
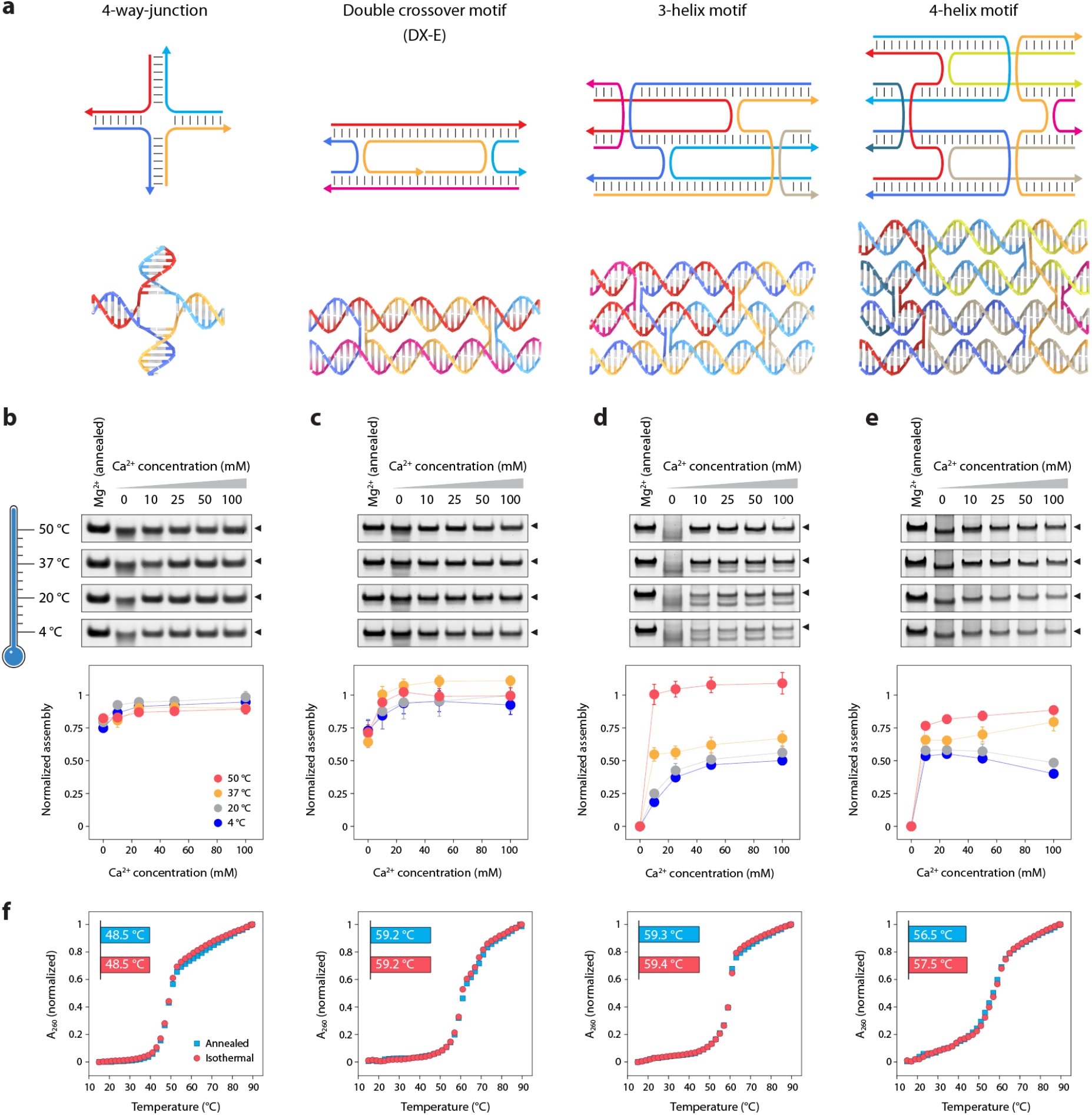
Isothermal assembly of DNA nanostructures in calcium. (a) Scheme and models of the DNA nanostructures used in this study. (b-e) Isothermal assembly at different temperatures (shown by the gradient) in buffer containing different Ca^2+^ concentrations for the 4-arm junction, DX-E, 3-helix motif, and 4-helix motif, respectively. (f) Melting curves of structures annealed and isothermally assembled in Ca^2+^. Data represent mean and error propagated from standard deviations of experiments performed in triplicates.

We then used an isothermally assembled tensegrity triangle motif for hierarchical assembly of designed three-dimensional (3D) DNA crystals.^34^ The tensegrity triangle motif contains three double-helical domains connected at the vertices by strand crossovers.^35^ The motif we use here contains three turns of DNA per edge (31 bp) and is three-fold symmetric; i.e. each duplex edge has the same sequence (**Fig. 4a, Supplementary Fig. 15, Supplementary Table 6**).^36^ The motif is tailed with sticky ends that allow them to cohere with each other to form a macroscopic 3D crystal (**Fig. 4b**). In prior research, authors of this work have created 3D DNA crystals using biologically-derived strands,^37^ studied the effect of sticky ends on crystal assembly and robustness,^38^ and used DNA crystals as a scaffold for triplex forming oligonucleotides.^39^ In the typical crystal assembly protocol, the component DNA strands are mixed together, set up in a hanging drop, and the entire multi-well plate taken through a thermal annealing protocol from 60-20 °C.^34,38^ In some cases, the motifs are annealed first and then set up for crystallization.^40^ Here, we annealed or isothermally assembled the motif at 50 °C in 25 mM Ca^2+^ and used the assembled sample for crystal setup. We observed rhombohedral crystals, the typical crystal habit observed for these motifs^34,37,38^, in both the annealed and isothermally assembled samples (**Fig. 4c**). Crystals from the isothermally assembled samples were 300-400 µm in almost all the hanging drops, while crystals obtained using the annealing protocol were generally smaller, with a few large crystals (**Fig. 4c** and **Supplementary Fig. 16**). These results show that isothermally assembled motifs can be used to create 3D DNA crystals, with potential use in encapsulating guests in a one-step process, determining ion-dependent response of crystals^41^ and enhancing thermal stability of DNA-nanoparticle colloidal crystals.^42^ There are still several aspects to be studied, such as the crystal growth rates, the effect of a combination of ions on crystal assembly, and the crystal structure of the designed lattice.

**Fig. 4.**
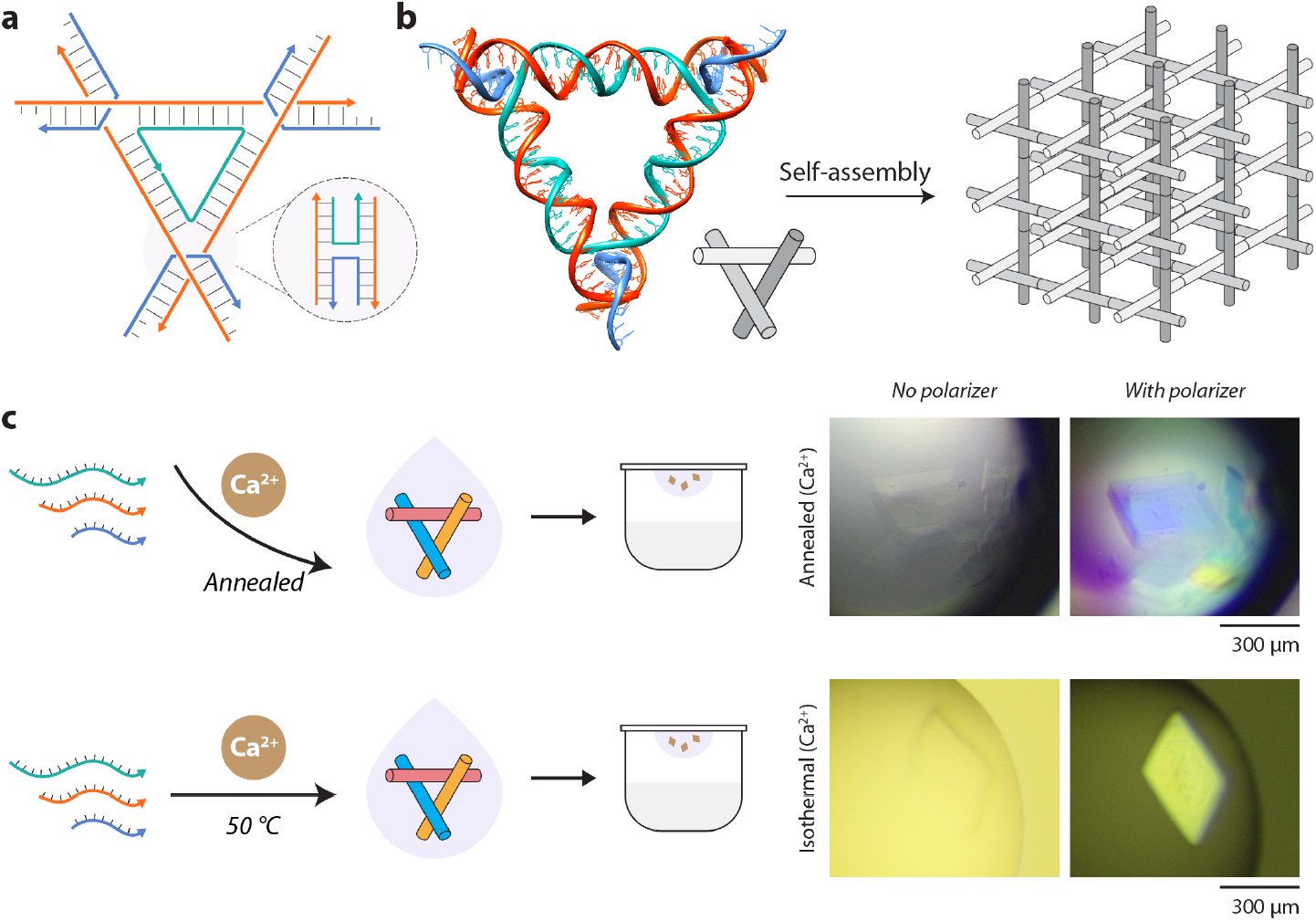
Rationally designed 3D DNA crystals assembled in calcium. (a) Scheme of the tensegrity triangle motif containing three double-helical edges connected by strand crossovers. (b) Model of the tensegrity triangle motif with 3 turns per edge and assembly into a crystal lattice. The triangle is 3-fold symmetric and contains the same sequence on each edge. The edges of the triangle are shown in different shades in the illustration for clarity. (c) Growth of 3D DNA crystals from motifs annealed or isothermally assembled in buffer containing calcium.

### Assembly and dynamic functionality of DNA tweezers in calcium

To demonstrate the isothermal assembly of a dynamic DNA nanostructure in Ca^2+^, we used a DNA tweezer^43^ that contains two double-helical domains connected by a single-stranded loop (open state). Each double helical domain contains a toehold that can bind a set strand S, resulting in the reconfiguration of the tweezer to the closed state (**Fig. 5a, Supplementary Fig. 17, Supplementary Table 7**). We first validated isothermal assembly of both the open and closed states of the DNA tweezer in 1× TAE-Ca^2+^ (**Supplementary Fig. 18**). Addition of the set strand S to the isothermally assembled DNA tweezer resulted in a 100% conversion of the open state to the closed state (**Fig. 5b and Supplementary Fig. 19**). We then added the unset strand S* to revert the tweezer back to the open state, followed by two more cycles of reconfiguration. The performance of isothermally assembled DNA tweezers with 10 mM Ca^2+^ was similar to that of the control sample prepared using a thermal annealing protocol in 10 mM Mg^2+^ (**Fig. 5c**).

**Fig. 5.**
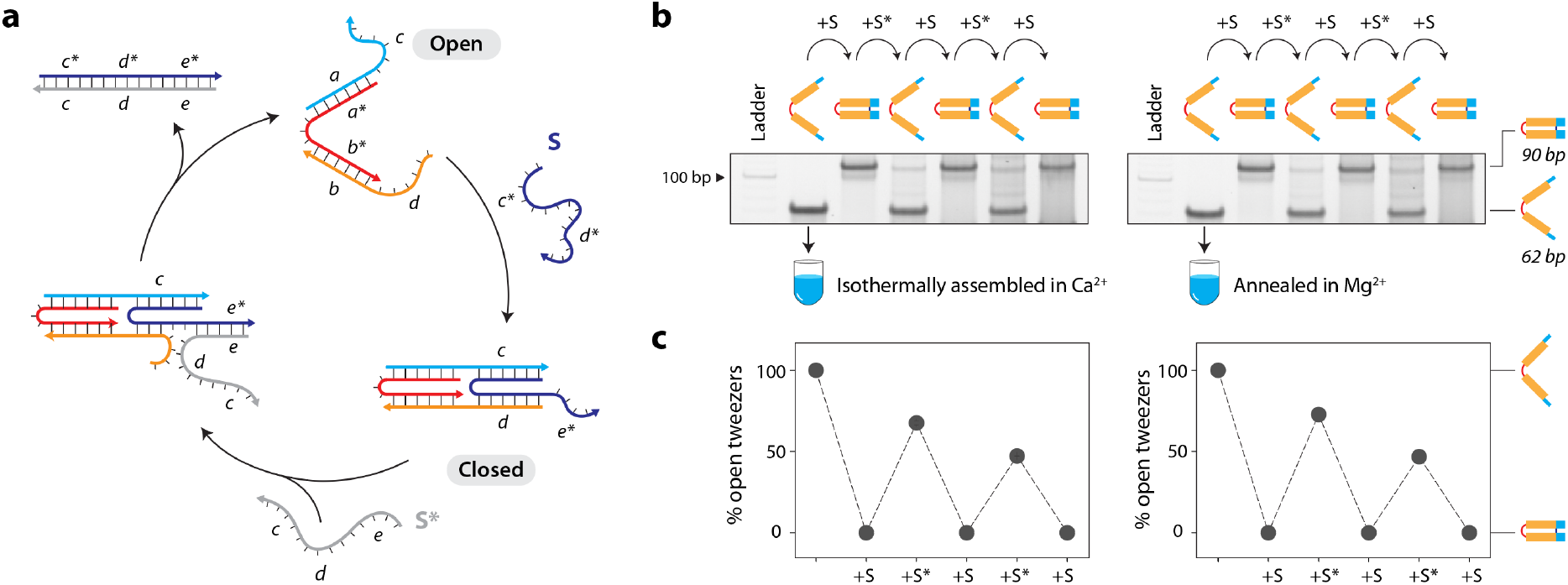
Assembly and functionality of DNA tweezers in calcium. (a) Design and operation of the DNA tweezer between the open and closed states by strand displacement. (b) Interconversion between open and closed states of the DNA tweezer starting with sample isothermally assembled in Ca^2+^ (left) or annealed in Mg^2+^ (right). (c) Conversion yields between open and closed states calculated from gels shown in (b). Data represent mean and error propagated from standard deviations of experiments performed in duplicates.

### Isothermal assembly in other ions

After demonstrating the successful isothermal assembly of various DNA motifs in a Ca^2+^-containing buffer, we tested the isothermal assembly of the DX-O motif in other metal ions. We assembled the DX-O motif in different concentrations of metal ions Li^+^, Na^+^, K^+^, Mg^2+^, Sr^2+^, Ni^2+^, Cu^2+^, Ag^+^, Zn^2+^, and Pb^2+^ isothermally at 4 °C, 20 °C, 37 °C, or 50 °C for 3 hours. We observed assembly of the DX-O motif in Na^+^, Li^+^, K^+^, Mg^2+^, and Sr^2+^ at all the temperatures tested, with an ion-concentration-dependent increase in assembly yields in most cases (**Fig. 6a-e, Supplementary Fig. 20**). In monovalent ions, the highest assembly yields were observed at 10 mM or 25 mM ion concentration at 37 °C. We found that the structure formed with very poor yields in Pb^2+^, Zn^2+^, Cu^2+^, and Ag^+^ at all the isothermal assembly temperatures tested, indicating degradation or aggregation of the DNA strands or structure (**Supplementary Fig. 21**). These ions are known to destabilize DNA duplexes^44^ and cause DNA cleavage,^45,46^ and have been known to affect the assembly of DNA nanostructures when using a thermal annealing process.^19^

**Fig. 6.**
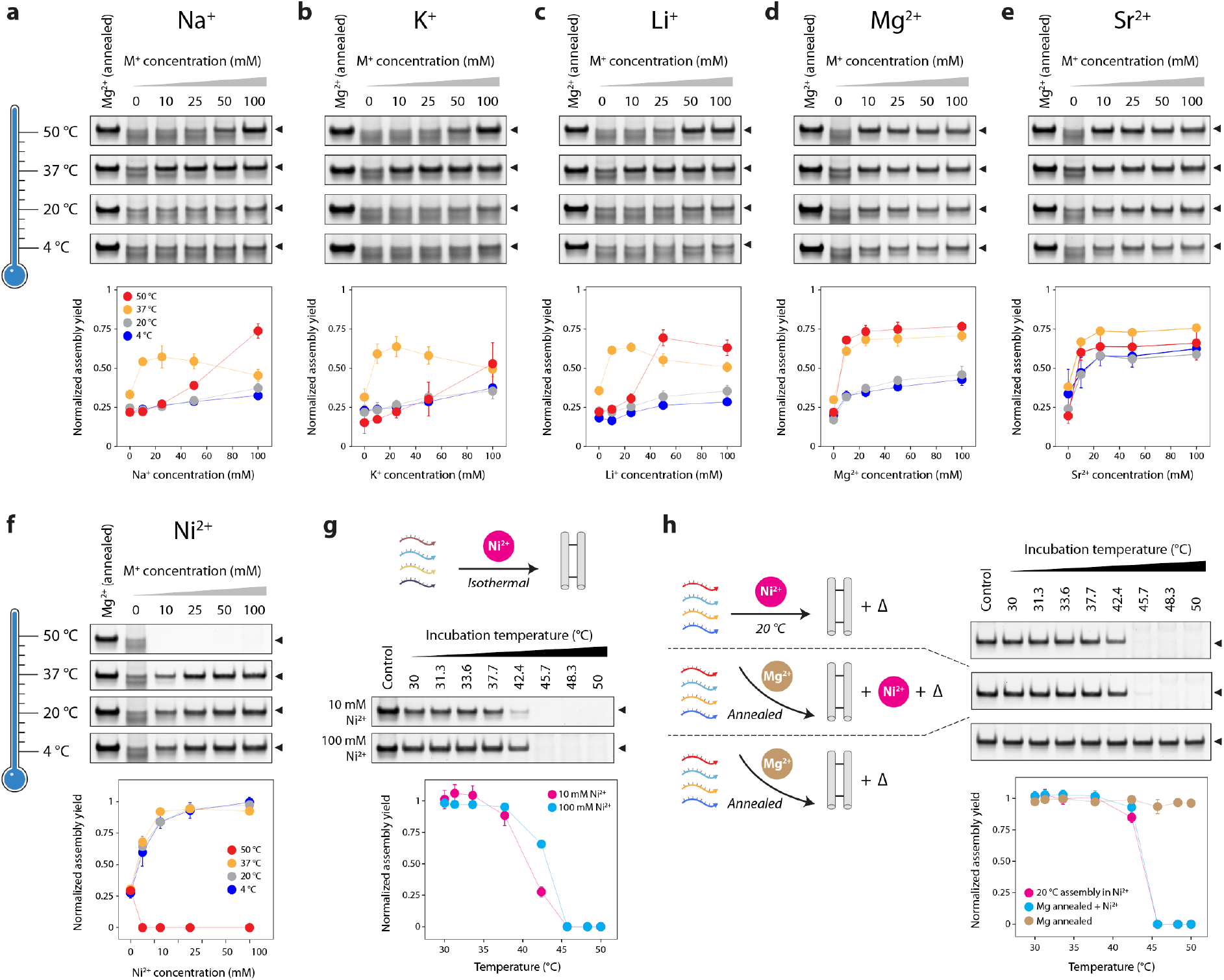
Isothermal assembly in different ions. (a-e) Isothermal assembly of DX-O motif in different ions at 4 °C, 20 °C, 37 °C and 50 °C. (f) Isothermal assembly of DX-O motif in Ni^+^ at 4 °C, 20 °C, 37 °C and 50 °C. (g) Assembly of DX-O in 10 and 100 mM Ni^2+^ incubated for 2 h at different temperatures from 30 to 50 °C. (h) Effect of Ni^2+^ and temperature on DX-O motif isothermally assembled in Ni^2+^ or annealed in Mg^2+^. Data represent mean and error propagated from standard deviations of experiments performed in triplicates.

In buffer containing Ni^2+^, we observed proper assembly of the DX-O motif when the strands were incubated at 4 °C, 20 °C, and 37 °C but not at 50 °C (**Fig. 6f, Supplementary Fig. 22**). This effect could be due to DNA damage or DNA cleavage caused by Ni^2+^ at higher temperatures.^46–48^ To identify the maximum viable isothermal assembly temperature for Ni^2+^, we incubated the DNA strand mixtures at a temperature range of 30-50 °C in 1× TAE containing 10 mM or 100 mM Ni^2+^ for 2 hours (**Fig. 6g, Supplementary Fig. 23**). We observed that assembly yields decreased with increasing temperature from 30-45 °C, with no assembly beyond this temperature. For DX-O motif isothermally assembled in Ni^2+^ at 20 °C and DX-O motif annealed in Mg^2+^ and then mixed with 100 mM Ni^2+^, we observed that the band corresponding to the structure was reduced or eliminated when incubated at temperatures above 40 °C post-assembly, indicating that Ni^2+^ causes degradation or aggregation of DNA structures at higher temperatures (**Fig. 6h and Supplementary Fig. 24**). In contrast, the DX-O motif annealed in Mg^2+^ or isothermally assembled in Ca^2+^ were not affected at the higher temperatures (**Fig. 6h and Supplementary Fig. 25**). Further, samples assembled isothermally at 20 °C in 100 mM Ni^2+^ were degraded within a few minutes at 50 °C, as were samples annealed in Mg^2+^ and mixed with 100 mM Ni^2+^ (**Supplementary Fig. 26**). This phenomenon was also observed in single-stranded DNA (**Supplementary Fig. 27**). Our results indicate that DNA degradation at 40 °C to 50 °C is only observed with Ni^2+^ and is not induced by common divalent alkali earth metal ions, and assembly challenges in Ni^2+^ can be overcome by isothermal assembly at temperatures below 40 °C.

### Molecular dynamics simulation of DX DNA motif in different counter ions

To understand the interaction of different counter ions with DNA nanostructures, we performed all-atom molecular dynamics (MD) simulations of the DX-O motif in select divalent (Mg^2+^, Ca^2+^) and monovalent (Na^+^, K^+^) ions The structure was simulated for 100 ns at 300 K, at a fixed ion concentration of 10 mM. To analyze the structural changes induced by each ion, we aligned the backbone of the dominant conformation of the motif in each case to the initial structure of the DX-O motif (**Fig. 7a**). The dominant conformation, representing the most prevalent conformation, was identified by clustering the simulation snapshots based on root mean square deviation (RMSD) and finding the structure representing the largest cluster. We observed that the overall structure of the DX-O motif was preserved in the presence of each of the four ions, with some conformational differences. For divalent ions Mg^2+^ and Ca^2+^, we observed that the RMSD of the structure did not deviate significantly from the initial structure over time, whereas the deviation was higher for monovalent ions (**Fig. 7b**). These results were consistent with a previous study that simulated a 512 bp section of DNA origami and showed large deviations from the initial structure in environments with only Na^+^ whereas there was minimal deviation in Mg^2+^.^49^ To further understand the local dynamics and flexibility, we calculated the per-residue root mean square fluctuations (RMSF) to assess the changes associated with individual nucleotides in different ions (**Fig. 7c**). We observed that a higher number of nucleotides in each strand showed higher RMSF values in the presence of monovalent ions compared to divalent ions (**Supplementary Table 8**). While the central core of the motif remained stable, the two “forks” at the ends exhibited increased opening and fluctuation with monovalent ions compared to divalent ions at the same 10 mM ion concentration. This suggests that monovalent ions result in more dynamic and less constrained conformations at the terminal regions of the motif. The conformational flexibility induced by the monovalent ions can potentially create new binding regions for other molecules (small molecule drugs or proteins, for example) that are unavailable in the absence of such flexibility. In contrast, the more rigid structure observed with divalent ions implies that these ions provide enhanced stability and play a critical role in maintaining the integrity of the DNA structure, allowing for more controlled interactions under similar ionic conditions.

**Fig. 7.**
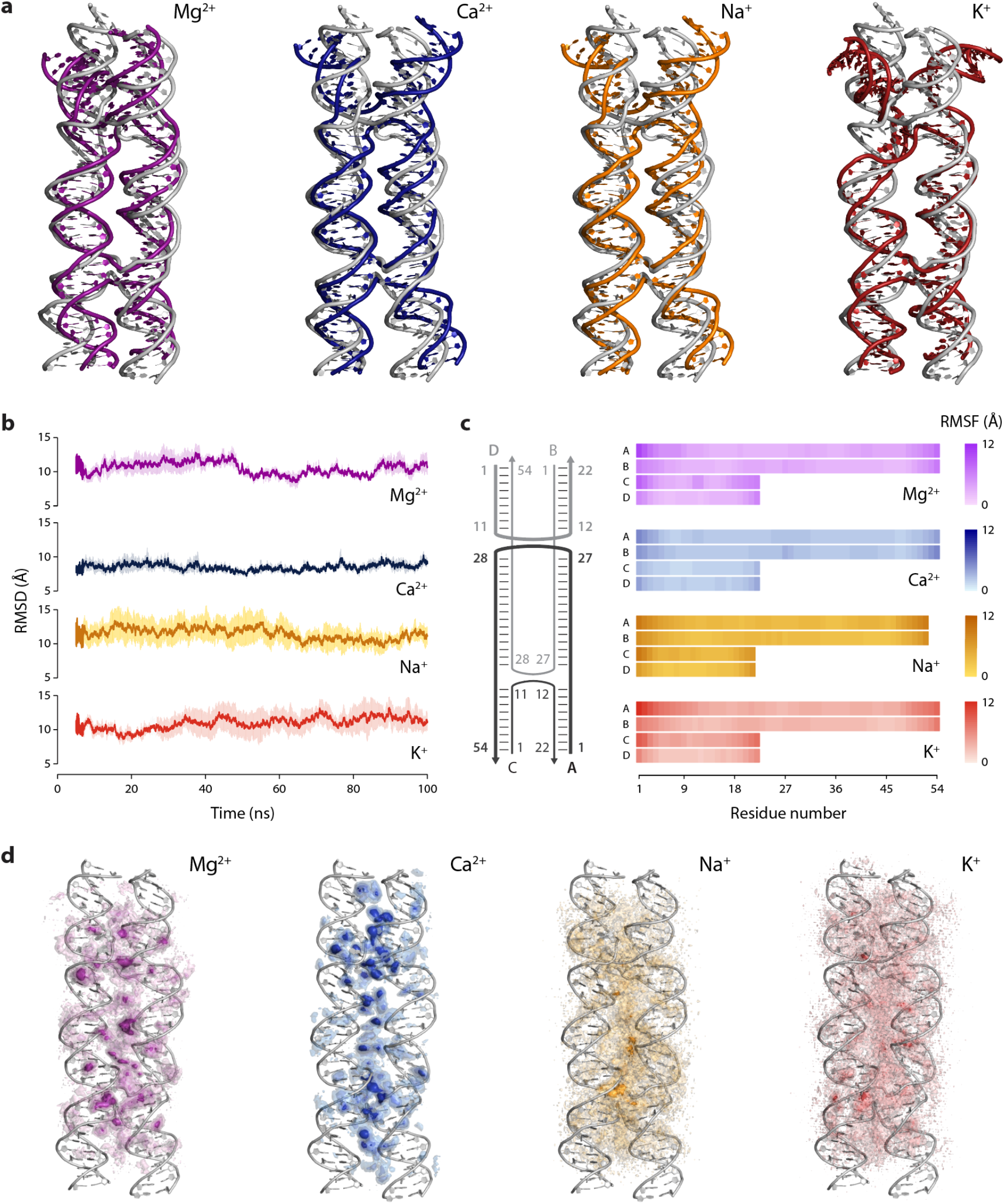
MD simulation of double crossover DNA motif in different cations. (a) Structure of the dominant conformation superimposed on the initial structure. (b) Root Mean Square Deviation (RMSD) of the DX-O motif. (c) Root Mean Square Fluctuation (RMSF) of the nucleotides in each strand of the motif. (d) Ion density around the motif shown as isomolarity surfaces that represent the time-averaged measure of the number of ions per unit volume in the vicinity of the DNA. Data shown in (b) to (d) are averages from three replicate 100 ns simulations.

To understand the ion distribution and binding preferences of the different ions, we calculated the ion density around the DNA, which is a time-averaged measure of the number of ions per unit volume in the vicinity of the DNA during the simulation (**Fig. 7d**). We observed that ions are more localized in the junctions and core of the motif where the backbones of multiple strands are closer together. The ion density was higher and more localized for divalent ions (Ca^2+^and Mg^2+^), suggesting stronger interactions and specific binding sites. Among the divalent ions, Ca^2+^interacted with DNA via stronger binding sites compared to Mg^2+^. In contrast, the ion density for both monovalent ions was more diffused, indicating less specific binding sites. Overall, our MD simulation results show that the interaction of Ca^2+^ ions with the DX-O motif is similar to that of Mg^2+^, with Ca^2+^ exhibiting a slightly higher stabilizing effect, while monovalent ions Na^+^ and K^+^ interact differently with the structure. This trend has been previously observed using ion exchange and spectrometric measurements,^50^ where the order of ion selectivity of DNA was determined to be Ca^2+^ > Mg^2+^ > K^+^ ≅ Na^+^. While the conformation of the motif is maintained in the presence of both monovalent and divalent ions, the latter provides greater structural stability at the same ion concentration.

### Biological feasibility of structures isothermally assembled in different cations

As DNA nanostructures are being used in biological applications, we then tested whether DNA motifs isothermally assembled in different ions interfere with cellular viability and immune response. The double crossover motif is a good model system for testing biological feasibility as it has been used for drug delivery applications.^51,52^ In previous work, we showed that DNA motifs assembled in Mg^2+^ do not affect the viability of HeLa cells^53^ and murine myoblast cells^54^ and do not interfere with the differentiation of myoblasts into myotubes.^54^ For this work, we incubated patient-derived fibroblasts, myoblasts, and myotubes with 500 nM DX-O isothermally assembled in different ions and performed the PrestoBlue cell viability assay. We did not observe any significant changes in cellular viability for any of these cell lines after 24 h of incubation with DX-O when compared to the viability of untreated cells (**Fig. 8a-c**). This observation was also similar to cells incubated with DX-O annealed in Mg^2+^. To delineate the effect of the different ions on cell viability, we incubated the cells with TAE buffer containing different cations (without any DNA). Our results confirmed that the different ions used for isothermal assembly did not significantly affect the viability of patient-derived fibroblasts, myoblasts, and myotubes at the concentrations tested here (**Supplementary Fig. 28**). Optical microscopy images of the cells incubated with DX-O showed no significant effect on cell appearance compared to untreated cells (**Fig. 8d** and **Supplementary Fig. 29-35**).

**Fig. 8.**
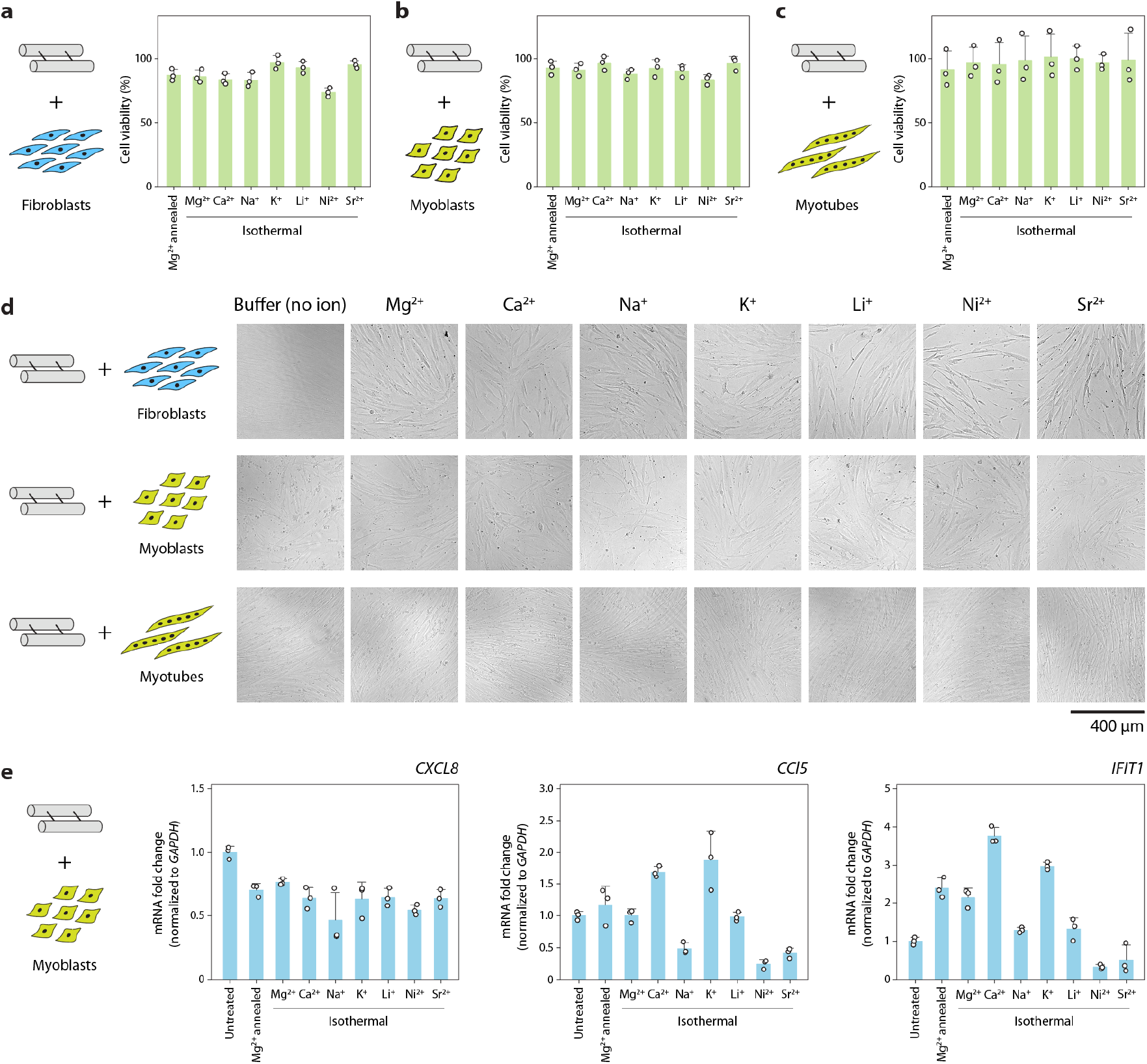
Effect of structures assembled isothermally in different ions on cell viability and immune response. Viability of (a) fibroblasts, (b) myoblasts, and (c) myotubes treated with DX-O motif isothermally assembled in different cations. Data is normalized to viability of untreated cells. (d) Brightfield microscopy images of cells treated with DX-O motif isothermally assembled in different cations for 24 h. (e) RT-qPCR analysis of immune response markers *CXCL8, CCL5, and IFIT1* in myoblasts incubated with DX-O motif isothermally assembled in different cations. Data represent mean and standard deviation from three biological replicates.

Next, we tested the immune response of cells to the DX-O motif isothermally assembled in different ions. We incubated myoblasts with 500 nM DX-O isothermally assembled in different ions for 24 h, isolated cellular RNA and performed RT-qPCR for three immune response markers: CXC motif chemokine ligand 8 (*CXCL-8*), chemokine (C-C motif) ligand 5 (*CCL5*), and interferon-induced protein with tetratricopeptide repeats 1 (*IFIT1*). Compared to untreated cells, we observed that the expression level of *CXCL-8* was marginally lower in all the ions used for assembly (**Fig. 8e**). For the other two markers tested here, we observed some changes in the immune response between different ions. Our recent work also showed that a nanostructured DNA motif induced lower expression of *CXCL-8* compared to duplex DNA,^53^ while other works have shown that the morphology and concentration of the nanostructure can determine their immune activation.^55^ Our findings show the potential feasibility of using DNA motifs assembled in different ions for biological applications.

## Discussion

In this work, we demonstrated the isothermal assembly of a variety of DNA motifs in different counterions. Although it is often overlooked for nanostructure assembly, Ca^2+^ is an effective substitute for Mg^2+^. As a counter ion of choice for isothermal assembly, Ca^2+^ is as effective as Mg^2+^ at generating 3D crystal lattices of tensegrity triangles and at producing dynamic devices such as DNA tweezers. We achieved over 75% assembly yield for the 4-arm junction, double crossover motif, 3-helix motif, and 4-helix motif under isothermal conditions at 50 °C. Simpler motifs such as 4-arm junction and double crossover motifs assembled efficiently at Ca^2+^ concentrations as low as 10 mM and at a temperature as low as 37 °C, while more complex 3-helix and 4-helix motifs required relatively higher Ca^2+^ concentration and temperature for their assembly. Similar to the role of temperature observed here, prior work showed that a lower starting temperature of 50 °C for thermal annealing decreased the folding efficiency to 50%,^56^ and isothermal assembly at 60 °C showed a higher hybridization rate and fewer defects compared to room temperature,^18^ but only studied assembly in Mg^2+^. Our work shows that in some cases, isothermal assembly allows rapid assembly of DNA motifs in contrast to standard annealing protocols. It remains to be seen how the kinetics of assembly varies for intra-strand folding (such as in DNA origami, where the staple strands connect different regions of the scaffold strand) and intermolecular assembly, where multiple DNA strands or component units associate to form a structure. Our study showed that compared to monovalent ions, divalent ions are a better choice as counter ions for the isothermal assembly of DX-O, as relatively higher concentrations and temperatures were needed to achieve over 75% yield when using monovalent counter-ions. These results are further supported by MD simulations that highlight the distinct modes of ion distribution patterns around the DX-O motif for monovalent and divalent ions. The influence of different ions on DNA self-assembly is also a key factor and combinations of different ions on isothermal DNA nanostructure assembly are yet to be tested. For example, in our previous study, we observed anomalous electrophoretic behavior of Ba^2+^-assembled structures^57^ (and therefore avoided in this study), and another study showed that a combination of metal ions protects DNA at high temperatures.^58^ In assessing different cations, our results indicate that for assembly in Ni^2+^, temperature-dependent depurination and strand cleavage of DNA^46–48^ can pose a problem for standard annealing methods, but can be overcome by isothermal assembly at lower temperatures. The facile nature of isothermal assembly could allow DNA nanostructures to be used as templates for synthesizing nickel nanostructures, such as nanowires, with applications in advanced materials and protein assembly.^59^

Isothermal assembly of DNA nanostructures is a key feature that can overcome the need for multi-step synthesis of protein-DNA hybrid nanostructures^7–9^ or the use of steep temperature gradients^60^ in annealing protocols to avoid heat damage to component molecules. Isothermal assembly of such finite DNA nanostructures indicates the possibility of using these structures for creating higher-order structures and arrays at constant temperatures using hybridization chain reaction and T-junctions,^61^ as well as the integration of strand displacement circuits and DNAzyme reactions for enhanced functionality.^62^ Isothermal assembly of nucleic acid strands into complex nanostructures, particularly using biologically relevant Ca^2+^ and Mg^2+^ ions, can be useful to fabricate functional nanodevices within living systems where a temperature gradient is unachievable.^63,64^ Such synthetic strategies can also be useful for DNA nanostructure-templated biomineralization.^65^ Overall, our work indicates the ability to assemble DNA nanostructures in physiological conditions for biological and materials science applications that require a constant environmental temperature and compatibility with several different counter ions.

## Acknowledgments

Research reported in this publication was supported by the National Institutes of Health (NIH) through the National Institute of General Medical Sciences (NIGMS) under award number R35GM150672 to A.R.C. and R35GM124720 to K.H., the National Institute on Aging (NIA) under award number R03AG076599 to A.R.C., and the National Institute of Neurological Disorders and Stroke (NINDS) under award number R01NS135254 to J.A.B and S.V. This work was also supported by the University at Albany Faculty Research Awards Programs (FRAP) seed funding to A.R.C., the USAMRAA CDMRP HT9425-24-1-0191 PRMRP to J.A.B and S.V, and Muscular Dystrophy Association award number 1192211 to J.A.B. A.R. was supported by the NIA Research Supplement to Promote Diversity in Health-Related Research awarded to A.R.C. (award R03AG076599). We thank John Cleary for comments on the manuscript.

## Author contributions

A.R., B.R.M., J.M., H.T., and V.M. performed assembly and characterization experiments. B.R.M. and A.R.C. designed experiments and analyzed the assembly and characterization data. K.S. performed cell studies and analyzed the corresponding data. Z.N. and S.V. performed molecular dynamics simulations and analyzed the corresponding data. J.A.B. supervised cell studies. A.R.C. conceived and supervised the project and visualized the data. B.R.M. and A.R.C. wrote the manuscript with edits from S.V., K.H. and J.A.B.

## Data availability

The data that support the findings of this study are available within the paper and its supplementary information files, including uncropped gel images of representative replicates for all experiments.

## References

1. Xavier, P. L. & Chandrasekaran, A. R. DNA-based construction at the nanoscale: emerging trends and applications. Nanotechnology 29, 062001 (2018).

2. Nickels, P. C. et al. Molecular force spectroscopy with a DNA origami–based nanoscopic force clamp. Science 354, 305–307 (2016).

3. Kilchherr, F. et al. Single-molecule dissection of stacking forces in DNA. Science 353, aaf5508 (2016).

4. Douglas, S. M., Chou, J. J. & Shih, W. M. DNA-nanotube-induced alignment of membrane proteins for NMR structure determination. PNAS 104, 6644–6648 (2007).

5. Khoshouei, A. et al. Designing Rigid DNA Origami Templates for Molecular Visualization Using Cryo-EM. Nano Lett. 24, 5031–5038 (2024).

6. Lee, H. et al. Molecularly self-assembled nucleic acid nanoparticles for targeted in vivo siRNA delivery. Nature Nanotechnology 7, 389–393 (2012).

7. Chandrasekaran, A. R. et al. DNA Nanoswitch Barcodes for Multiplexed Biomarker Profiling. Nano Lett. 21, 469–475 (2021).

8. Zhang, C. et al. Symmetry Controls the Face Geometry of DNA Polyhedra. J. Am. Chem. Soc. 131, 1413–1415 (2009).

9. Flory, J. D. et al. PNA-Peptide Assembly in a 3D DNA Nanocage at Room Temperature. J. Am. Chem. Soc. 135, 6985–6993 (2013).

10. Sobczak, J.-P. J., Martin, T. G., Gerling, T. & Dietz, H. Rapid Folding of DNA into Nanoscale Shapes at Constant Temperature. Science 338, 1458–1461 (2012).

11. Myhrvold, C., Dai, M., Silver, P. A. & Yin, P. Isothermal Self-Assembly of Complex DNA Structures under Diverse and Biocompatible Conditions. Nano Lett. 13, 4242–4248 (2013).

12. Shi, X., Zhao, H., Li, X. & Song, T. Isothermal approach to assemble spatial DNA nanotubes for drug delivery. Oncotarget 5, (2018).

13. Rossi-Gendron, C. et al. Isothermal self-assembly of multicomponent and evolutive DNA nanostructures. Nat. Nanotechnol. 18, 1311–1318 (2023).

14. Jungmann, R., Liedl, T., Sobey, T. L., Shih, W. & Simmel, F. C. Isothermal Assembly of DNA Origami Structures Using Denaturing Agents. J. Am. Chem. Soc. 130, 10062–10063 (2008).

15. Zhang, Z., Song, J., Besenbacher, F., Dong, M. & Gothelf, K. V. Self-Assembly of DNA Origami and Single-Stranded Tile Structures at Room Temperature. Angewandte Chemie International Edition 52, 9219–9223 (2013).

16. Kopielski, A., Schneider, A., Csáki, A. & Fritzsche, W. Isothermal DNA origami folding: avoiding denaturing conditions for one-pot, hybrid-component annealing. Nanoscale 7, 2102–2106 (2015).

17. Ji, B. et al. Room Temperature Study of Seeding Growth on Two-Dimensional DNA Nanostructure. Langmuir 35, 4140–4145 (2019).

18. Song, J. et al. Isothermal Hybridization Kinetics of DNA Assembly of Two-Dimensional DNA Origami. Small 9, 2954–2959 (2013).

19. Rodriguez, A. et al. Self-Assembly of DNA Nanostructures in Different Cations. Small 19, 2300040 (2023).

20. Zhou, K. et al. Boosted Productivity in Single-Tile-Based DNA Polyhedra Assembly by Simple Cation Replacement. ChemBioChem 23, e202200138 (2022).

21. Chou, L. Y. T., Song, F. & Chan, W. C. W. Engineering the Structure and Properties of DNA-Nanoparticle Superstructures Using Polyvalent Counterions. J. Am. Chem. Soc. 138, 4565–4572 (2016).

22. Kielar, C. et al. On the Stability of DNA Origami Nanostructures in Low-Magnesium Buffers. Angewandte Chemie International Edition 57, 9470–9474 (2018).

23. Martin, T. G. & Dietz, H. Magnesium-free self-assembly of multi-layer DNA objects. Nat Commun 3, 1103 (2012).

24. Wang, D. et al. Isothermal Self-Assembly of Spermidine–DNA Nanostructure Complex as a Functional Platform for Cancer Therapy. ACS Appl. Mater. Interfaces 10, 15504–15516 (2018).

25. Li, Y. et al. Universal pH-Responsive and Metal-Ion-Free Self-Assembly of DNA Nanostructures. Angewandte Chemie International Edition 57, 6892–6895 (2018).

26. Fu, T. J. & Seeman, N. C. DNA double-crossover molecules. Biochemistry 32, 3211–3220 (1993).

27. Kallenbach, N. R., Ma, R.-I. & Seeman, N. C. An immobile nucleic acid junction constructed from oligonucleotides. Nature 305, 829–831 (1983).

28. LaBean, T. H. et al. Construction, Analysis, Ligation, and Self-Assembly of DNA Triple Crossover Complexes. J. Am. Chem. Soc. 122, 1848–1860 (2000).

29. Reishus, D., Shaw, B., Brun, Y., Chelyapov, N. & Adleman, L. Self-Assembly of DNA Double-Double Crossover Complexes into High-Density, Doubly Connected, Planar Structures. J. Am. Chem. Soc. 127, 17590–17591 (2005).

30. Wang, R., Kuzuya, A., Liu, W. & Seeman, N. C. Blunt-ended DNA stacking interactions in a 3-helix motif. Chem. Commun. 46, 4905–4907 (2010).

31. Liu, D., Park, S. H., Reif, J. H. & LaBean, T. H. DNA nanotubes self-assembled from triple-crossover tiles as templates for conductive nanowires. Proceedings of the National Academy of Sciences 101, 717–722 (2004).

32. Mao, C., LaBean, T. H., Reif, J. H. & Seeman, N. C. Logical computation using algorithmic self-assembly of DNA triple-crossover molecules. Nature 407, 493–496 (2000).

33. Bardales, A. C., Mills, J. R. & Kolpashchikov, D. M. DNA Nanostructures as Catalysts: Double Crossover Tile-Assisted 5′ to 5′ and 3′ to 3′ Chemical Ligation of Oligonucleotides. Bioconjugate Chem. 35, 28–33 (2024).

34. Zheng, J. et al. From molecular to macroscopic via the rational design of a self-assembled 3D DNA crystal. Nature 461, 74–77 (2009).

35. Liu, D., Wang, M., Deng, Z., Walulu, R. & Mao, C. Tensegrity: Construction of Rigid DNA Triangles with Flexible Four-Arm DNA Junctions. J. Am. Chem. Soc. 126, 2324–2325 (2004).

36. Nguyen, N. et al. The absence of tertiary interactions in a self-assembled DNA crystal structure. Journal of Molecular Recognition 25, 494–494 (2012).

37. Sha, R. et al. Self-Assembled DNA Crystals: The Impact on Resolution of 5′-Phosphates and the DNA Source. Nano Lett. 13, 793–797 (2013).

38. Ohayon, Y. P. et al. Designing Higher Resolution Self-Assembled 3D DNA Crystals via Strand Terminus Modifications. ACS Nano 13, 7957–7965 (2019).

39. Rusling, D. A. et al. Functionalizing Designer DNA Crystals with a Triple-Helical Veneer. Angewandte Chemie International Edition 53, 3979–3982 (2014).

40. Li, Z. et al. Making Engineered 3D DNA Crystals Robust. J. Am. Chem. Soc. 141, 15850–15855 (2019).

41. Zheng, M., Li, Z., Zhang, C., Seeman, N. C. & Mao, C. Powering ≈50 µm Motion by a Molecular Event in DNA Crystals. Advanced Materials 34, 2200441 (2022).

42. Calcaterra, H. A., Chellam, N. S., Lee, B., Schatz, G. C. & Mirkin, C. A. High Temperature, Isothermal Growth Promotes Close Packing and Thermal Stability in DNA-Engineered Colloidal Crystals. ACS Nano (2024) doi:10.1021/acsnano.4c09308.

43. Yurke, B., Turberfield, A. J., Mills Jr, A. P., Simmel, F. C. & Neumann, J. L. A DNA-fuelled molecular machine made of DNA. Nature 406, 605–608 (2000).

44. Morris, D. L. DNA-bound metal ions: recent developments. Biomolecular Concepts 5, 397–407 (2014).

45. Rodríguez, M. R. et al. DNA cleavage mechanism by metal complexes of Cu(II), Zn(II) and VO(IV) with a schiff-base ligand. Biochimie 186, 43–50 (2021).

46. Guo, H. et al. Nickel Carcinogenesis Mechanism: DNA Damage. International Journal of Molecular Sciences 20, 4690 (2019).

47. Robison, S. H., Cantoni, O. & Costa, M. Strand breakage and decreased molecular weight of DNA induced by specific metal compounds. Carcinogenesis 3, 657–662 (1982).

48. Nackerdien, Z., Kasprzak, K. S., Rao, G., Halliwell, B. & Dizdaroglu, M. Nickel(II)- and Cobalt(II)-dependent Damage by Hydrogen Peroxide to the DNA Bases in Isolated Human Chromatin1. Cancer Research 51, 5837–5842 (1991).

49. Roodhuizen, J. A. L., Hendrikx, P. J. T. M., Hilbers, P. A. J., de Greef, T. F. A. & Markvoort, A. J. Counterion-Dependent Mechanisms of DNA Origami Nanostructure Stabilization Revealed by Atomistic Molecular Simulation. ACS Nano 13, 10798–10809 (2019).

50. Korolev, N., Lyubartsev, A. P., Rupprecht, A. & Nordenskiöld, L. Competitive Binding of Mg2+, Ca2+, Na+, and K+ Ions to DNA in Oriented DNA Fibers: Experimental and Monte Carlo Simulation Results. Biophysical Journal 77, 2736–2749 (1999).

51. Stewart, J. M. et al. Programmable RNA microstructures for coordinated delivery of siRNAs. Nanoscale 8, 17542–17550 (2016).

52. Zhang, H. et al. DNA nanostructures coordinate gene silencing in mature plants. PNAS 116, 7543–7548 (2019).

53. Madhanagopal, B. R. et al. The unusual structural properties and potential biological relevance of switchback DNA. Nat Commun 15, 6636 (2024).

54. Chandrasekaran, A. R. et al. Exceptional Nuclease Resistance of Paranemic Crossover (PX) DNA and Crossover-Dependent Biostability of DNA Motifs. J. Am. Chem. Soc. 142, 6814–6821 (2020).

55. Hu, Y. et al. Morphology Dictated Immune Activation with Framework Nucleic Acids. Small 19, 2303454 (2023).

56. Kuzyk, A., Laitinen, K. T. & Törmä, P. DNA origami as a nanoscale template for protein assembly. Nanotechnology 20, 235305 (2009).

57. Madhanagopal, B. R., Rodriguez, A., Cordones, M. & Chandrasekaran, A. R. Barium Concentration-Dependent Anomalous Electrophoresis of Synthetic DNA Motifs. ACS Appl. Bio Mater. 7, 2704–2709 (2024).

58. Lu, C. et al. Protection of DNA by metal ions at 95 °C: from lower critical solution temperature (LCST) behavior to coordination-driven self-assembly. Nanoscale 14, 14613–14622 (2022).

59. Becerril, H. A., Ludtke, P., Willardson, B. M. & Woolley, A. T. DNA-Templated Nickel Nanostructures and Protein Assemblies. Langmuir 22, 10140–10144 (2006).

60. Fu, J. et al. Multi-enzyme complexes on DNA scaffolds capable of substrate channelling with an artificial swinging arm. Nature Nanotechnology 9, 531–536 (2014).

61. Nie, Z., Wang, P., Tian, C. & Mao, C. Synchronization of Two Assembly Processes To Build Responsive DNA Nanostructures. Angewandte Chemie International Edition 53, 8402–8405 (2014).

62. Zhang, D. Y., Hariadi, R. F., Choi, H. M. T. & Winfree, E. Integrating DNA strand-displacement circuitry with DNA tile self-assembly. Nat Commun 4, 1965 (2013).

63. Lin, C. et al. In vivo cloning of artificial DNA nanostructures. PNAS 105, 17626–17631 (2008).

64. Li, M. et al. In vivo production of RNA nanostructures via programmed folding of single-stranded RNAs. Nature Communications 9, 2196 (2018).

65. Athanasiadou, D. & Carneiro, K. M. M. DNA nanostructures as templates for biomineralization. Nat Rev Chem 5, 93–108 (2021).

