## Supplementary information for "Influence of metal ions on the isothermal self-assembly of DNA nanostructures"

### Materials and methods

#### Preparation of DNA complexes

DNA strands were purchased from Integrated DNA Technologies (IDT). For different metal ions, the chemicals used were lithium chloride (LiCl, Sigma Aldrich), sodium chloride (NaCl, Sigma-Aldrich), potassium chloride (KCl, Sigma-Aldrich), magnesium chloride ($MgCl_2$, Sigma-Aldrich), calcium chloride dihydrate ($CaCl_2·2H_2O$, Sigma-Aldrich), strontium chloride ($SrCl_2$, Sigma-Aldrich), zinc chloride anhydrous ($ZnCl_2$, Sterm chemicals), silver nitrate, ($AgNO_3$, Millipore), lead (II) chloride ($PbCl_2$, Sigma-Aldrich), nickel chloride hexahydrate, ($NiCl_2·6H_2O$, VWR). For control DNA complexes annealed in $Mg^{2+}$, component DNA strands were mixed in 1× tris-acetate EDTA (1× TAE) buffer containing 40 mM Tris base (pH 8.0), 20 mM acetic acid, 2 mM EDTA containing 12.5 mM $Mg^{2+}$. For samples annealed in other ions, component DNA strands were mixed in 1× TAE containing the specified amounts of the corresponding metal salt. Samples were annealed in a thermal cycler with the following steps: 90 °C for 3 minutes, 65 °C for 20 minutes, 45 °C for 20 minutes, 37 °C for 30 minutes, 20 °C for 30 minutes, and the solution was then cooled to 4°C. For isothermal assembly, component DNA strands were mixed in 1× TAE containing the specified amounts of metal salt and incubated at 4°C, 20 °C, 37 °C, or 50 °C for specific time durations.

#### Non-denaturing polyacrylamide gel electrophoresis

DNA samples were mixed with 10× loading dye containing bromophenol blue and glycerol and loaded in polyacrylamide gels (19:1 acrylamide solution, National Diagnostics) prepared in 1× TAE buffer containing 12.5 mM $Mg^{2+}$. Gels were run at 4 °C with 1× TAE-$Mg^{2+}$ as the running buffer and stained with 0.5× GelRed (Biotium) in water for 20 min in dark and destained in water for 10 min. Gels were imaged on a Bio-Rad Gel Doc XR+ imager using the default settings for GelRed with UV illumination and analyzed using ImageLab software (Bio-Rad).

#### Circular dichroism (CD) spectroscopy

CD spectra were collected on 1 μM DNA complexes assembled in 1× TAE buffer containing $Mg^{2+}$ or $Ca^{2+}$ at the concentrations specified in the text. Experiments were performed on a Jasco-815 CD spectrometer at room temperature in a quartz cell with a 1 mm path length. CD spectra were collected from 360 to 200 nm with a scanning speed of 100 nm/min, bandwidth of 1 nm, and digital integration time was set at 1 s. The average of three accumulations were recorded.

#### Thermal melting studies

UV thermal melting studies were performed on DNA complexes assembled in 1× TAE containing 50 mM $Mg^{2+}$ or $Ca^{2+}$. The concentrations of DX-O, 4-arm junction, DX-E, 3-helix motif, and 4-helix motif were 1 μM, 2.38 μM, 1 μM, 0.69 μM, and 0.51 μM, respectively. Experiments were performed in a Cary 3500 UV-Visible Spectrophotometer equipped with a temperature controller. Melting curves were acquired at 260 nm by heating and cooling from 20 °C to 90 °C at a rate of 0.5 °C/min. The data were fitted to the Boltzmann function using OriginPro. Melting temperatures were determined from the first derivative of the fitted melting curves.

**Crystallization**

Crystals were grown in 5 µl hanging drops containing 5 µM DNA tensegrity triangle solution annealed or isothermally assembled in 1× TAE containing 25 mM $Ca^{2+}$. Drops were equilibrated against a 600 ml reservoir of 1.75 M ammonium sulfate at room temperature. The following protocol was used for annealed samples: 90 °C for 3 minutes, 65 °C for 20 minutes, 45 °C for 20 minutes, 37 °C for 30 minutes, 20 °C for 30 minutes. For isothermal assembly, samples were incubated at 50 °C for 6 h. Crystals were imaged using a Zeiss SteREO Discovery V12 microscope.

**Molecular dynamics simulations**

The initial all atom structure of the DX-O motif was modeled in Molecular Operating Environment (MOE).[1] This was achieved by generating two double stranded DNAs and introducing strand breaks and joined at the appropriate locations to create the DX-O motif. Molecular dynamics (MD) simulations were then performed using GROMACS 2019.4[2] to assess the stability and interaction of the structure with two monovalent ($Na^+$, $K^+$) and two divalent ($Mg^{2+}$, $Ca^{2+}$) ions. The amber99-bsc01 force-field was used for DNA.[3] The ion parameters reported in Schwartz et al.[4] were used for the monovalent and divalent ions. All simulations were carried out at 300 K and 10 mM ion concentration. Three replicate simulations of 100 ns were performed for the system.

The MD simulation incorporated a leap-frog algorithm with a 2-fs time step to integrate the equations of motion. The system was simulated using the velocity rescaling thermostat to maintain the temperature.[5] The pressure was maintained at 1 atm using the Berendsen barostat for equilibration.[6] Long-range electrostatic interactions were calculated using the particle mesh Ewald (PME) algorithm with a real space cut-off of 1.0 nm.[7] Lennard–Jones interactions were truncated at 1.0 nm. Water molecules were represented with the TIP3P model,[8] and the LINCS algorithm[9] was used to constrain the motion of hydrogen atoms bonded to heavy atoms. The system was subjected to energy minimization to prevent any overlap of atoms, followed by a 100 ns MD simulation. The first 5 ns of the trajectory were excluded from the analysis for system equilibration. Coordinates were stored every 2 ps for further analysis. The simulations were visualized using PyMOL.[10] The Root Mean Square Deviation (RMSD) and Root Mean Square Fluctuations (RMSF) were calculated using tools *rms* and *rmsf* in GROMACS.[2] The ion density around the DNA was calculated as ions per $Å^3$ using the density tool of the MDAnalysis module in python.[11] The density was converted to an equivalent molarity value and was visualized in PyMOL[10] as isomolarity surfaces which represent regions with the same ion density around the DNA.

**Cell culture, differentiation, and treatment**

Primary fibroblasts from an apparently healthy 37-year-old female individual (GM08400, control) were obtained from Corriell. Fibroblasts were cultured in 1× MEM (Corning) supplemented with 1×

antibiotic-antimycotic (Gibco) and 15% heat inactivated fetal bovine serum until they reached ~80-90% confluency. They were then seeded in 96 or 24 well plates. Once seeded, fibroblasts were grown to ≥90% confluency before beginning treatment with DNA nanostructures or buffers for 24 hours. A 5 μM solution of DX-O was prepared by incubating the DNA strands at 50 °C for 3 hours in buffers with different metal ions at 10 mM concentration and used for the cell studies.

Primary myoblasts derived from muscle biopsies of an apparently healthy 62-year-old female individual (150-05a, control) were obtained from Cell Applications. All myoblasts were grown in in SkGMTM-2 BulletKitTM growth medium (Lonza). Once the myoblasts reached ~80-90% confluency they were treated with either treated DNA nanostructures or buffers for 24 hours. To generate myotubes, myoblasts at ~80-90% confluency were differentiated by removing the Lonza medium and growing them in DMEM/F-12 50/50 medium (Corning) supplemented with 2% horse serum for 4 days. After differentiation myotubes were treated with DNA nanostructures or buffers for 24 hours.

**Cell viability assays**

Cells were seeded at ~$2x10^4$ cells/ml fibroblasts and myoblasts in 96 well plates and treated with DNA nanostructures or buffers. After treatment, samples were incubated with PrestoBlue™ HS Cell Viability Reagent (Thermo Scientific) for 1 hour at 37 °C and incubated in the dark. The absorbance at 570 and 600 nm was measured with a Synergy H1 plate reader. The 570/600 ratio was calculated for all wells and normalized to the DMSO treated cells. For myotubes, the cells were treated post-differentiation, and the same process was followed for viability assay.

**RT-qPCR transcript level analysis**

RNA was extracted using the NucleoSpin® 96 RNA extraction kit (Takarabio) following its manual which includes an in-column DNase I treatment. RNA concentrations of samples were measured with NanoDrop (Thermo Scientific). All RNA samples were reverse transcribed to make cDNA with SuperScript IV reverse transcriptase (Thermo Scientific) and random hexamers from IDT with a 30-minute incubation at 50 °C. The cDNA was used for RT-qPCR.

RT-qPCR was carried out using PowerUp SYBR (Applied Biosystems) according to manufacturer's instructions. Reactions were run on a the CFX384 Touch Real-Time PCR System (Bio-Rad) and analyzed on MS Excel using the 2-ΔΔCt method for gene expression. For myoblasts, the transcript levels of CXCL-8, CCL-5 and IFIT-1 were normalized to GAPDH. Relative transcript levels were calculated compared to untreated cells.

### References


1. Molecular Operating Environment (MOE), 2019.01; Chemical Computing Group ULC, 1010 Sherbrooke St. West, Suite #910, Montreal, QC, Canada, H3A 2R7, 2019.
2. Abraham, M. J. *et al.* GROMACS: High performance molecular simulations through multi-level parallelism from laptops to supercomputers. *SoftwareX* **1–2**, 19–25 (2015).
3. Ivani, I. *et al.* Parmbsc1: a refined force field for DNA simulations. *Nature Methods* **13**, 55–58 (2016).
4. Mamatkulov, S. & Schwierz, N. Force fields for monovalent and divalent metal cations in TIP3P water based on thermodynamic and kinetic properties. *The Journal of Chemical Physics* **148**, 074504 (2018).
5. Bussi, G., Donadio, D. & Parrinello, M. Canonical sampling through velocity rescaling. *J. Chem. Phys.* **126**, 014101 (2007).
6. Berendsen, H. J. C., Postma, J. P. M., van Gunsteren, W. F., DiNola, A. & Haak, J. R. Molecular dynamics with coupling to an external bath. *J. Chem. Phys.* **81**, 3684–3690 (1984).
7. Darden, T., York, D. & Pedersen, L. Particle mesh Ewald: An N·log(N) method for Ewald sums in large systems. *J. Chem. Phys.* **98**, 10089–10092 (1993).
8. Mark, P. & Nilsson, L. Structure and Dynamics of the TIP3P, SPC, and SPC/E Water Models at 298 K. *J. Phys. Chem. A* **105**, 9954–9960 (2001).
9. Hess, B., Bekker, H., Berendsen, H. J. C. & Fraaije, J. G. E. M. LINCS: A linear constraint solver for molecular simulations. *Journal of Computational Chemistry* **18**, 1463–1472 (1997).
10. Schrödinger, LLC. The PyMOL Molecular Graphics System, Version 1.8. (2015).
11. MDAnalysis: A Python Package for the Rapid Analysis of Molecular Dynamics Simulations - SciPy Proceedings. https://proceedings.scipy.org/articles/Majora-629e541a-00e (2016).

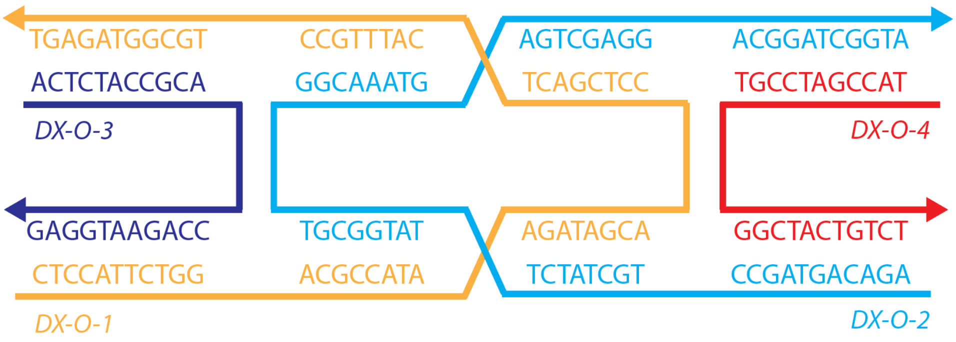


**Supplementary Figure 1**. Sequence design of the DX-O motif.

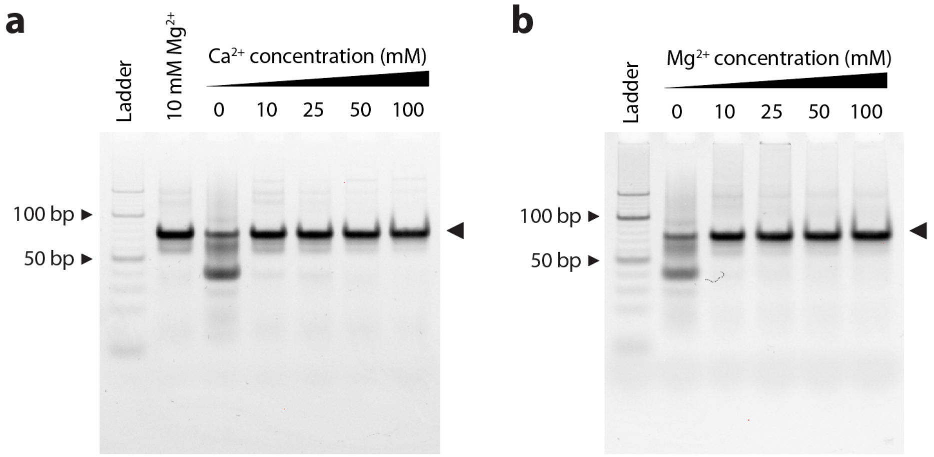


**Supplementary Figure 2**. (a) Non-denaturing polyacrylamide gel image of DX-O assembled in 1× TAE buffer containing different concentrations of $Ca^{2+}$ following a thermal annealing protocol. Lane 2 shows DX-O assembled in 10 mM $Mg^{2+}$ using the same annealing protocol. Full gel image of data shown in Fig 2c. (b) Non-denaturing polyacrylamide gel image of DX-O assembled in 1× TAE buffer containing different concentrations of $Mg^{2+}$ following a thermal annealing protocol. Data are normalized to assembly yield in 10 mM $Mg^{2+}$. Full gel image of data shown in Fig 2d.

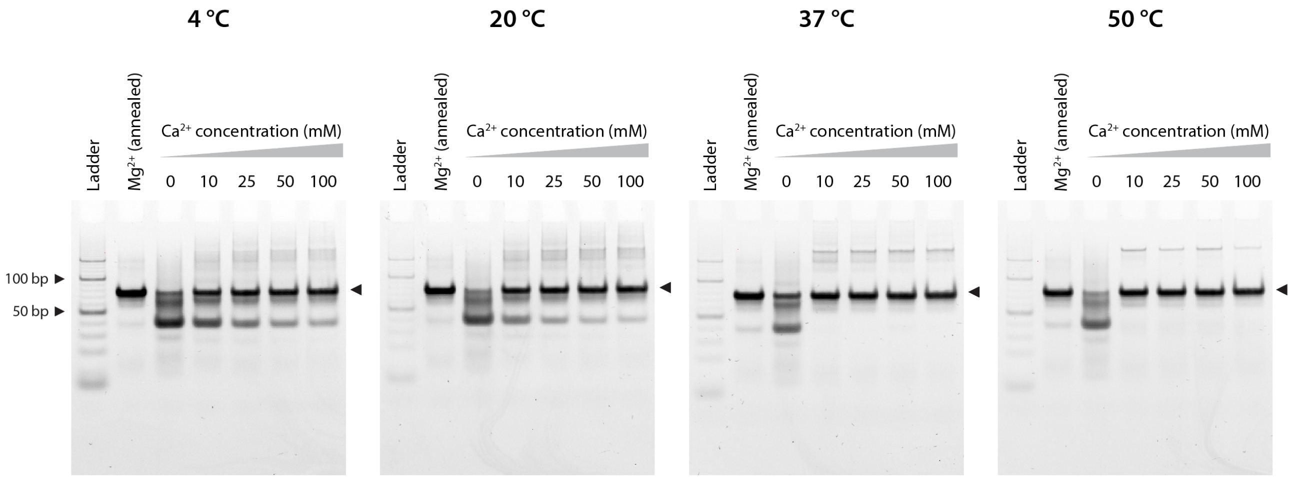


**Supplementary Figure 3**. Non-denaturing polyacrylamide gel images of DX-O assembled in 1× TAE buffer containing different concentrations of $Ca^{2+}$ following the isothermal protocol at different temperatures. Lane 2 in all the gels shows DX-O assembled in $Mg^{2+}$ using the annealing protocol. Full gel images of data shown in Fig 2e.

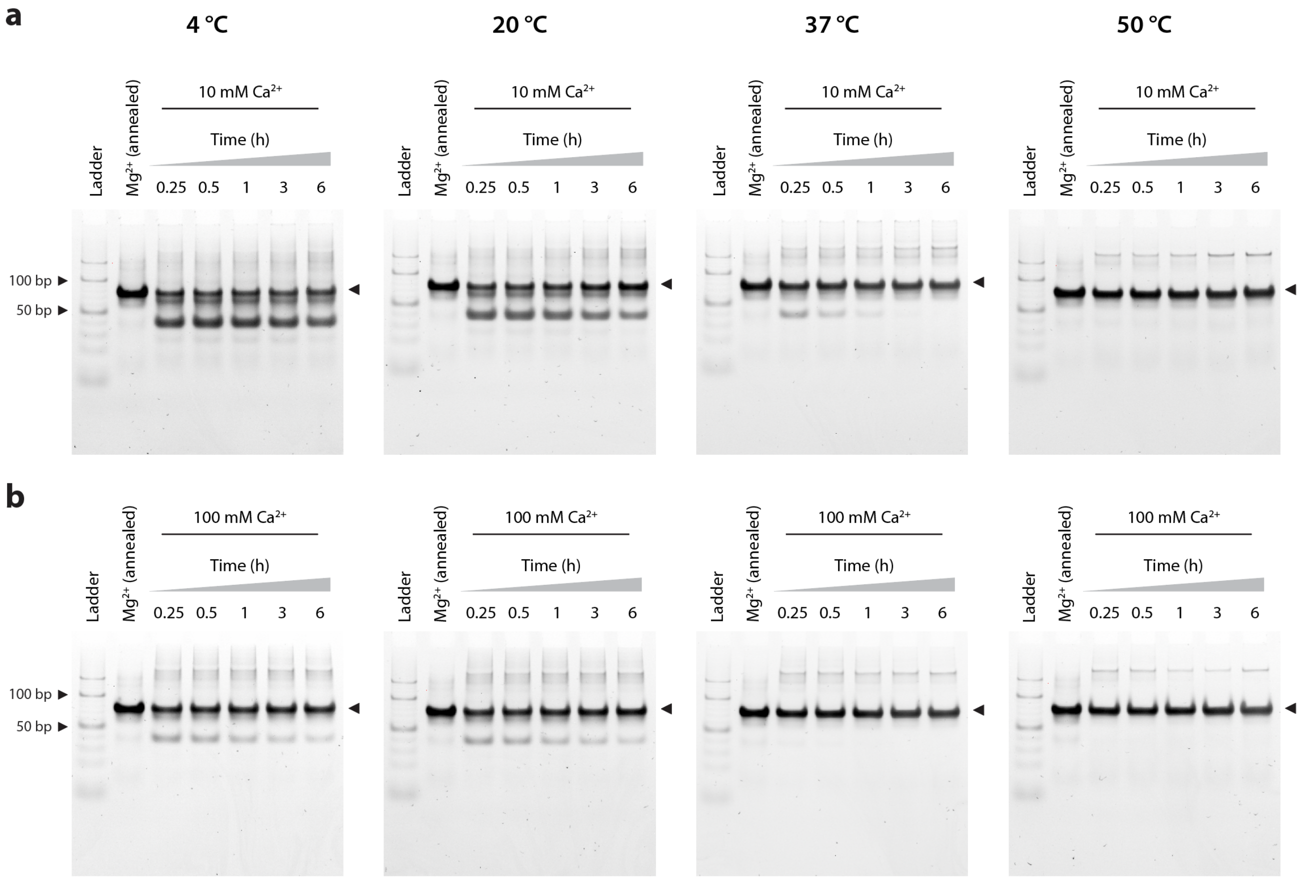


**Supplementary Figure 4**. Non-denaturing polyacrylamide gel images showing the assembly of DX-O under isothermal conditions in 1× TAE buffer containing (a) 10 mM $Ca^{2+}$ and (b) 100 mM $Ca^{2+}$, over different incubation periods and at different assembly temperatures. Lane 2 in all the gels shows DX-O assembled in $Mg^{2+}$ using the annealing protocol. (a) and (b) are full gel images of data shown in Fig 2f and 2g, respectively.

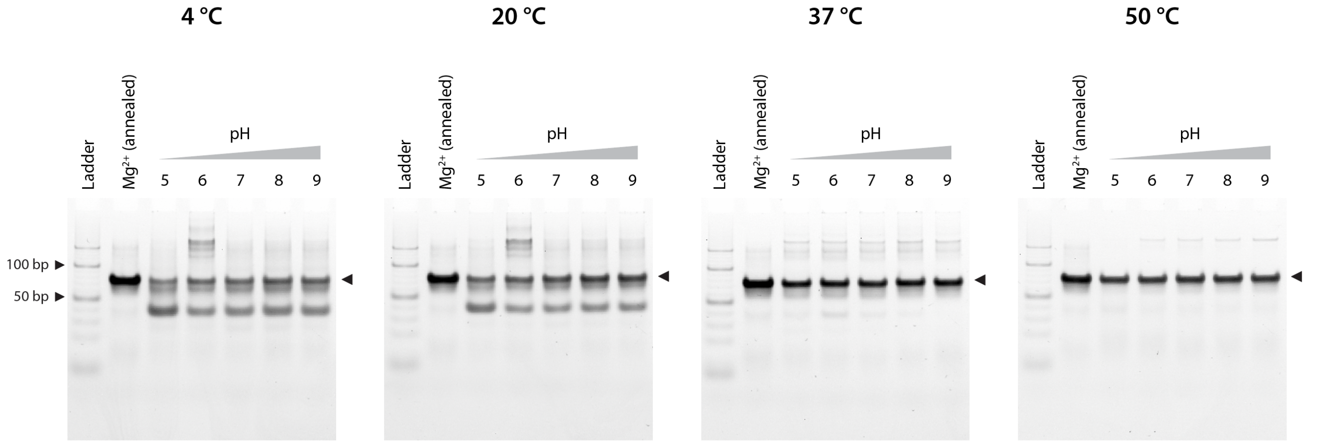


**Supplementary Figure 5**. Non-denaturing polyacrylamide gel images showing the assembly of DX-O under isothermal conditions in 1× TAE buffer at different pH and assembly temperatures. Lane 2 in all the gels shows DX-O assembled in $Mg^{2+}$ using the annealing protocol. Full gel images of data shown in Fig 2h.

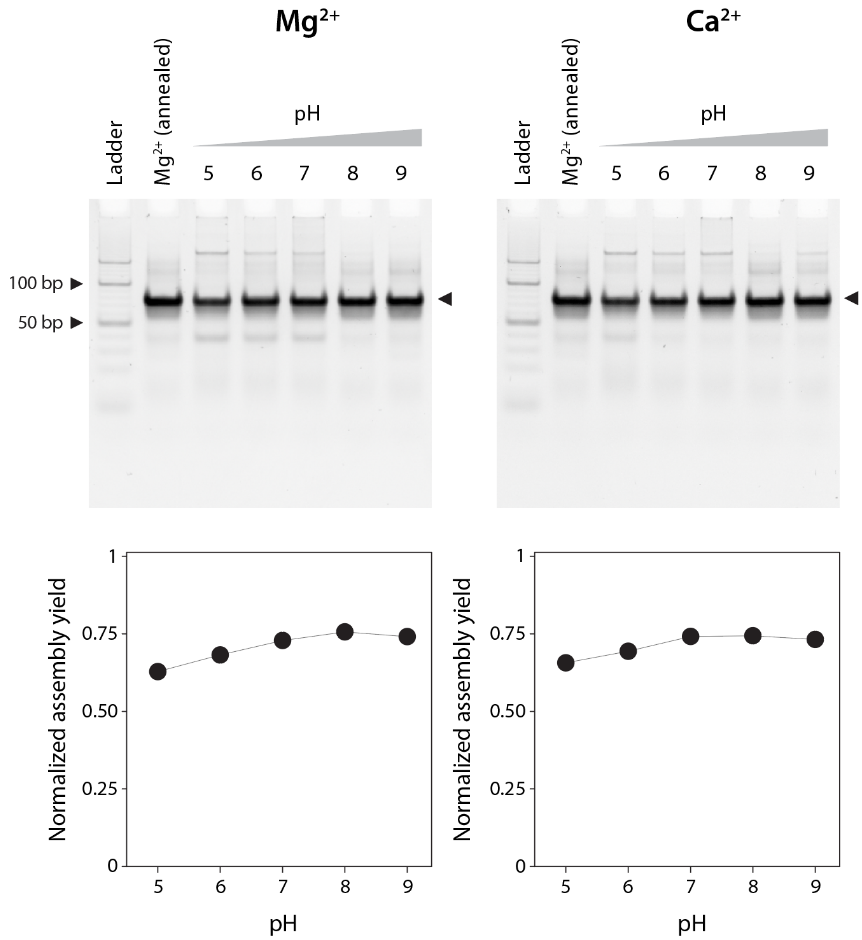


**Supplementary Figure 6**. Non-denaturing polyacrylamide gel images showing the yield of DX-O when assembled using the thermal annealing protocol in 1× TAE buffer containing $Mg^{2+}$ or $Ca^{2+}$ at different pH. Assembly yields are normalized to the assembly yield of the DX-O annealed in 1×TAE/$Mg^{2+}$ buffer at pH 8 (lane 2 in gels).

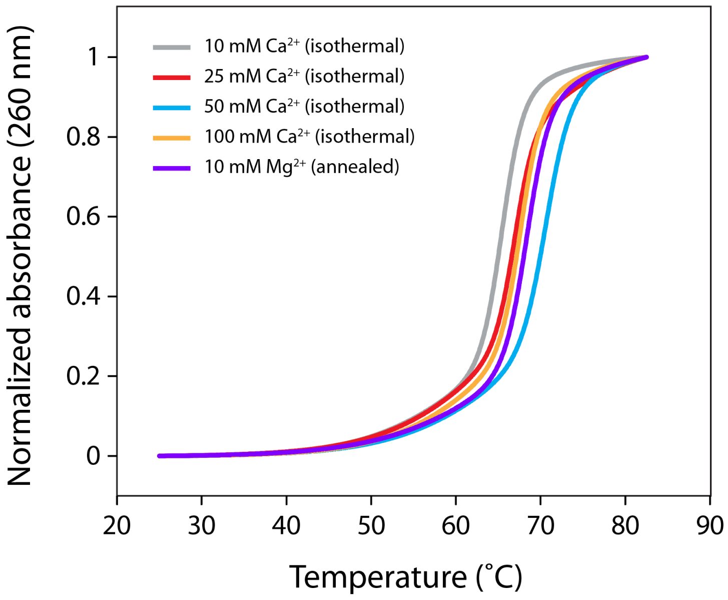


**Supplementary Figure 7**. Normalized UV-thermal melting curves of DX-O assembled in 1× TAE buffer containing different concentrations of $Ca^{2+}$ under the isothermal condition at 20 °C for 3 h and DX-O assembled in the buffer containing 10 mM $Mg^{2+}$ using the annealing protocol.

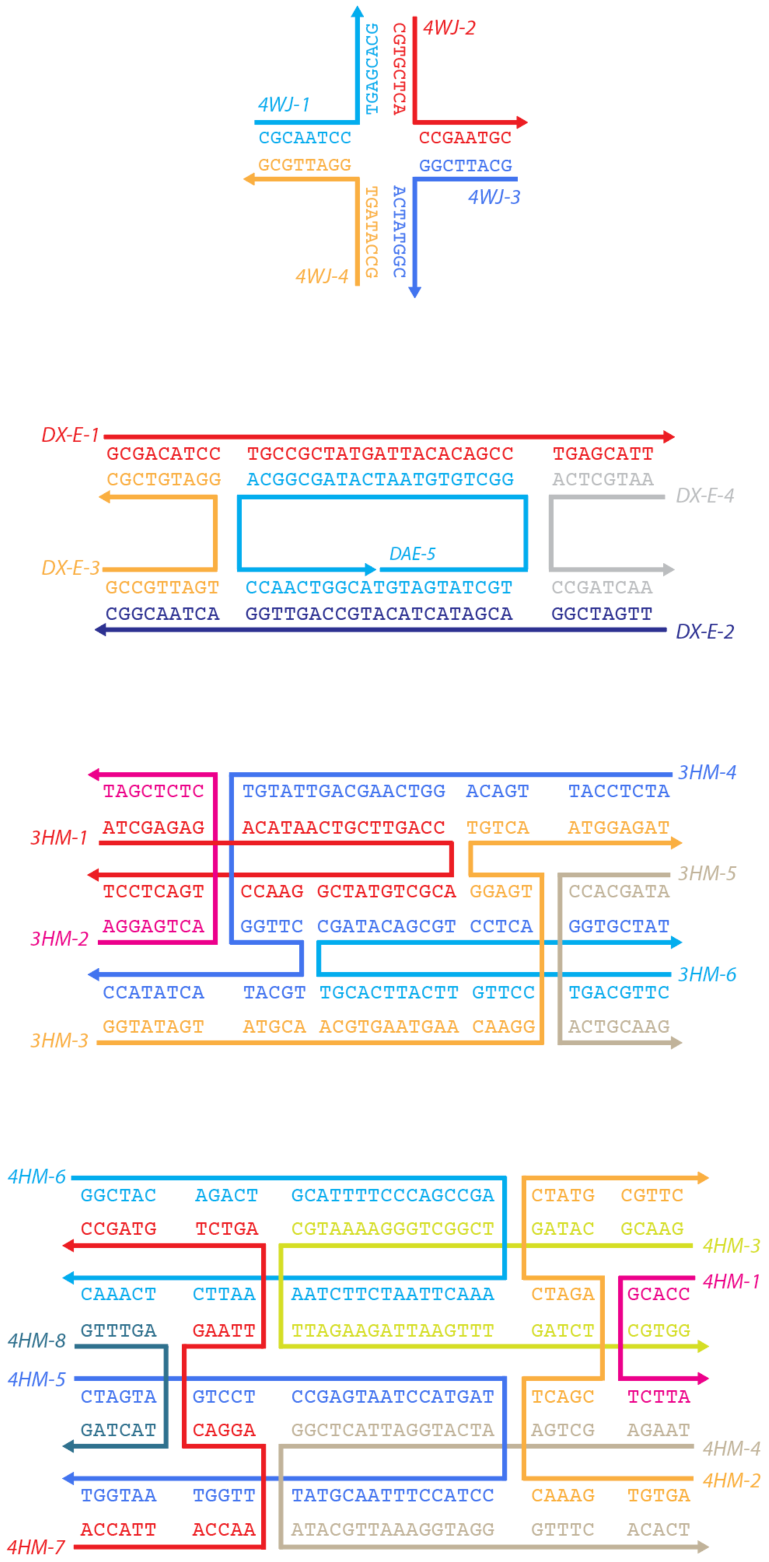


**Supplementary Figure 8**. Sequence and scheme of four-way junction, double crossover with even half-turns between two antiparallel crossovers (DX-E), 3-helix motif, and 4-helix motif.

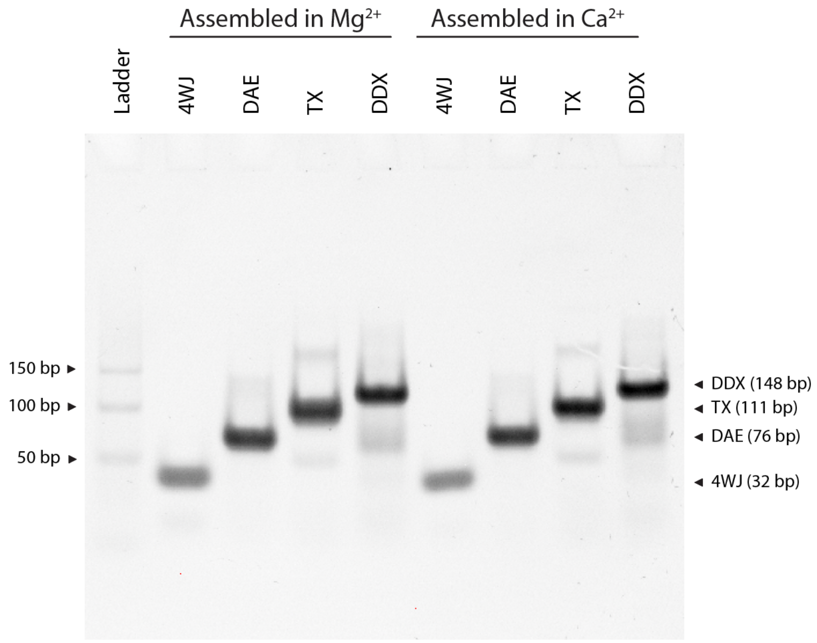


**Supplementary Figure 9**. Assembly of four-way junction, double crossover with even half-turns between two antiparallel crossovers (DX-E), 3-helix motif, and 4-helix motif using thermal annealing protocol in buffer containing 10 mM $Mg^{2+}$ or 10 mM $Ca^{2+}$.

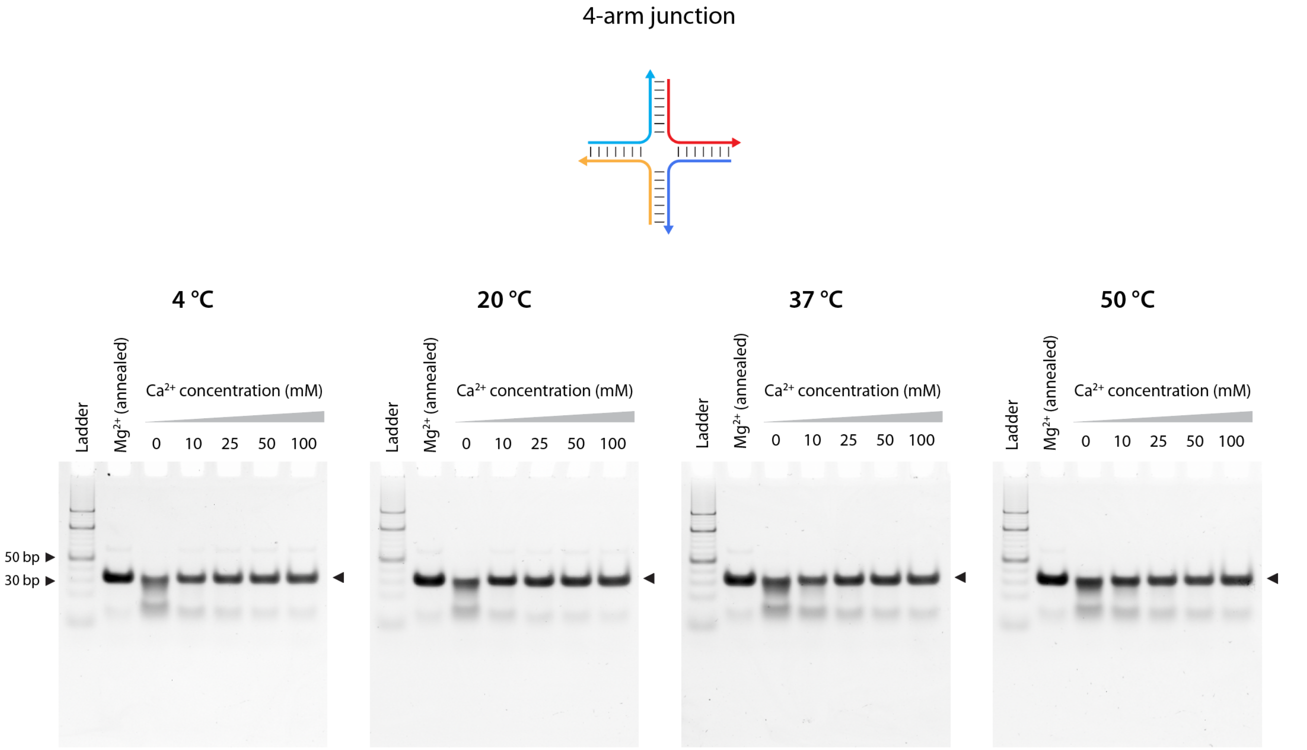


**Supplementary Figure 10**. Non-denaturing polyacrylamide gel images of four-way junction assembled in 1× TAE buffer containing different concentrations of $Ca^{2+}$ following the isothermal protocol at different temperatures. Lane 2 in all the gels shows four-way junction assembled in $Mg^{2+}$ using the annealing protocol. Full gels of the images in Fig 3b.

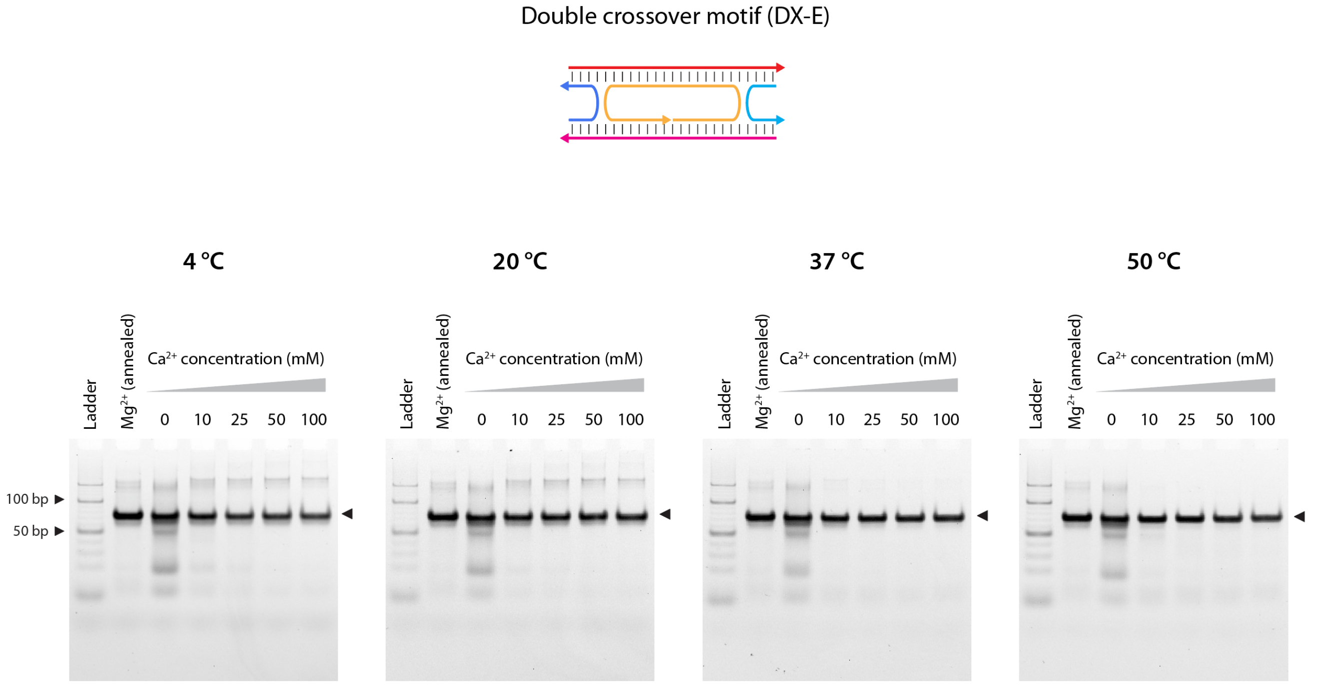


**Supplementary Figure 11**. Non-denaturing polyacrylamide gel images of double crossover (DX-E) assembled in 1× TAE buffer containing different concentrations of $Ca^{2+}$ following the isothermal protocol at different temperatures. Lane 2 in all the gels shows DX-E assembled in $Mg^{2+}$ using the annealing protocol. Full gels of the images in Fig 3c.

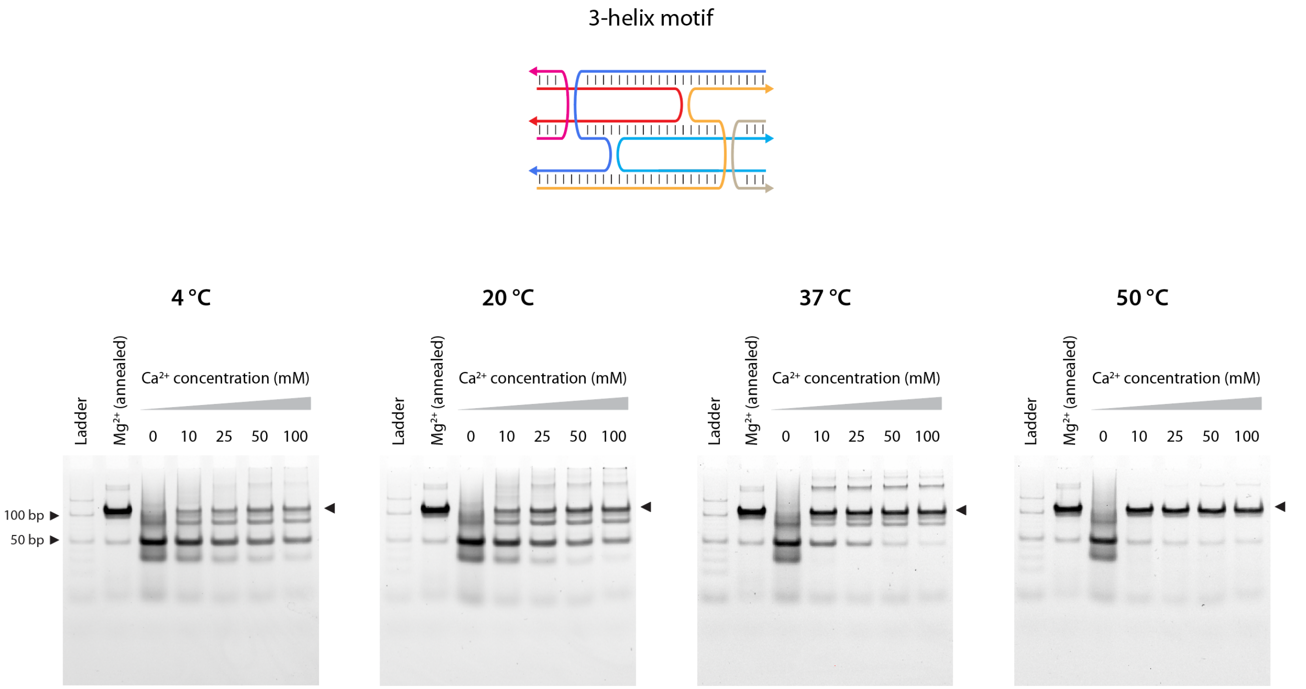


**Supplementary Figure 12**. Non-denaturing polyacrylamide gel images of 3-helix motif assembled in 1× TAE buffer containing different concentrations of $Ca^{2+}$ following the isothermal protocol at different temperatures. Lane 2 in all the gels shows 3-helix motif assembled in $Mg^{2+}$ using the annealing protocol. Full gels of the images in Fig 3d.

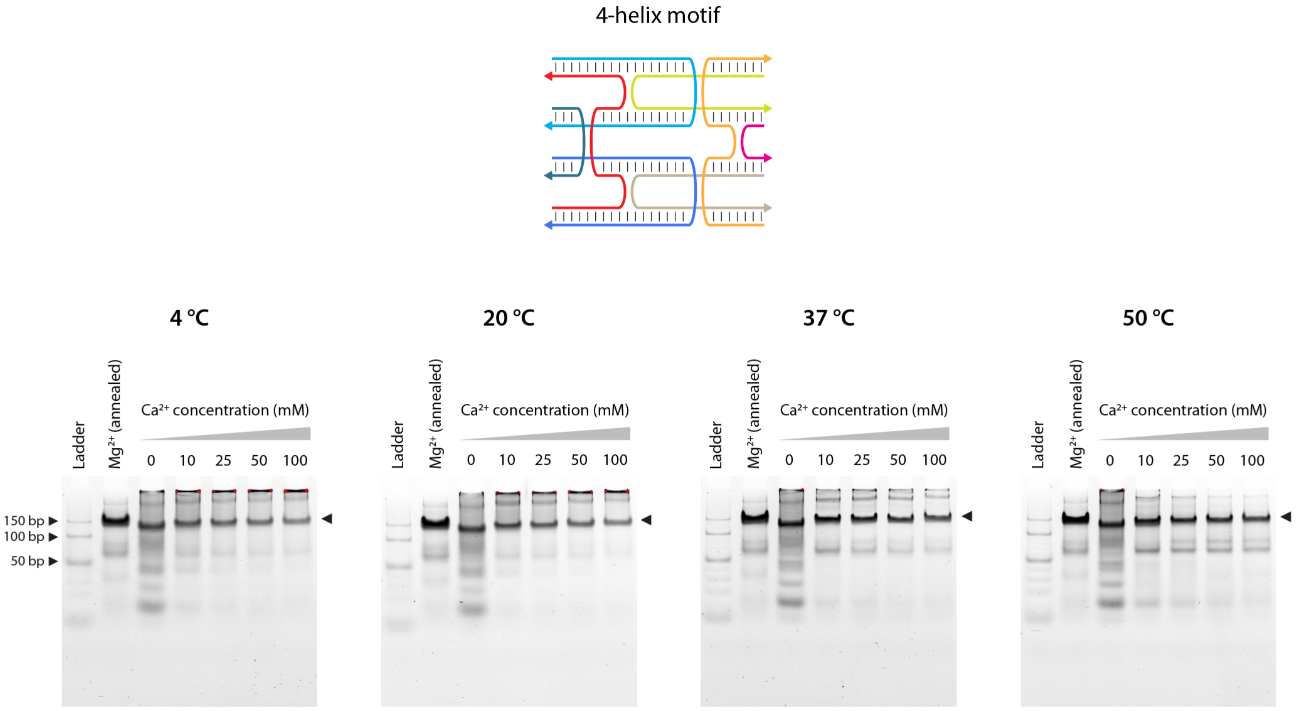


**Supplementary Figure 13**. Non-denaturing polyacrylamide gel images of 4-helix motif assembled in 1× TAE buffer containing different concentrations of $Ca^{2+}$ following the isothermal protocol at different temperatures. Lane 2 in all the gels shows 4-helix motif assembled in $Mg^{2+}$ using the annealing protocol. Full gels of the images in Fig 3e.

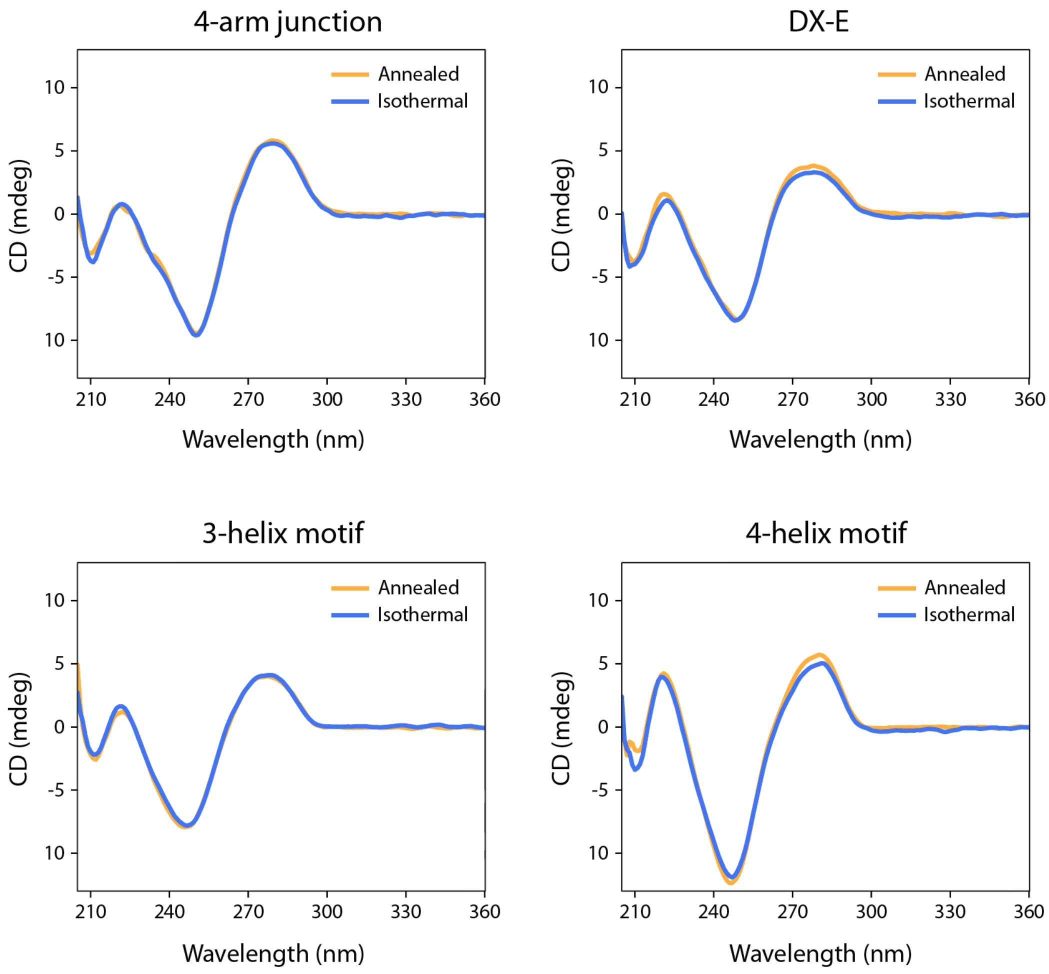


**Supplementary Figure 14**. Circular dichroism curves of 4-arm junction, double crossover with even half-turns between two antiparallel crossovers (DX-E), 3-helix motif, and 4-helix motif assembled in 1× TAE buffer containing different concentrations of $Ca^{2+}$ under the isothermal condition at 50 °C or using the annealing protocol.

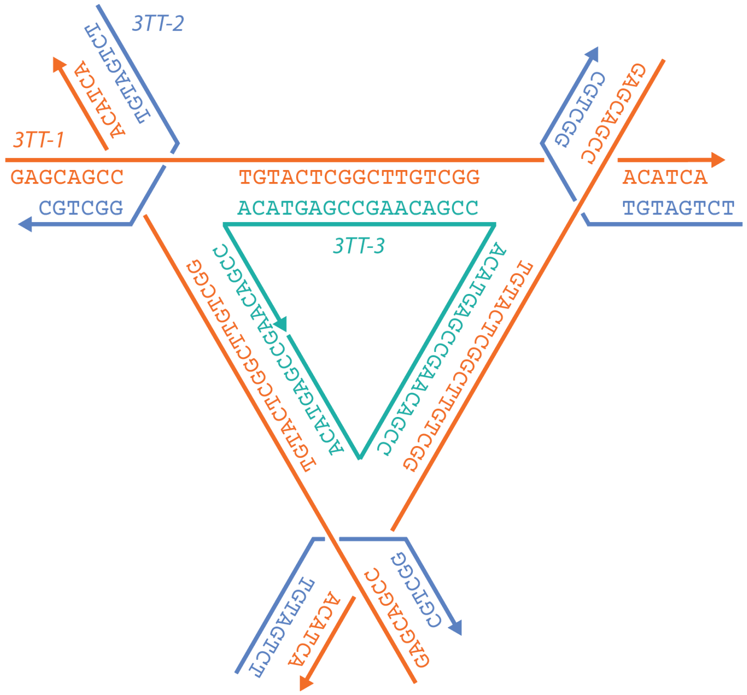


**Supplementary Figure 15**. Sequence and scheme of the 3-turn tensegrity triangle DNA motif.

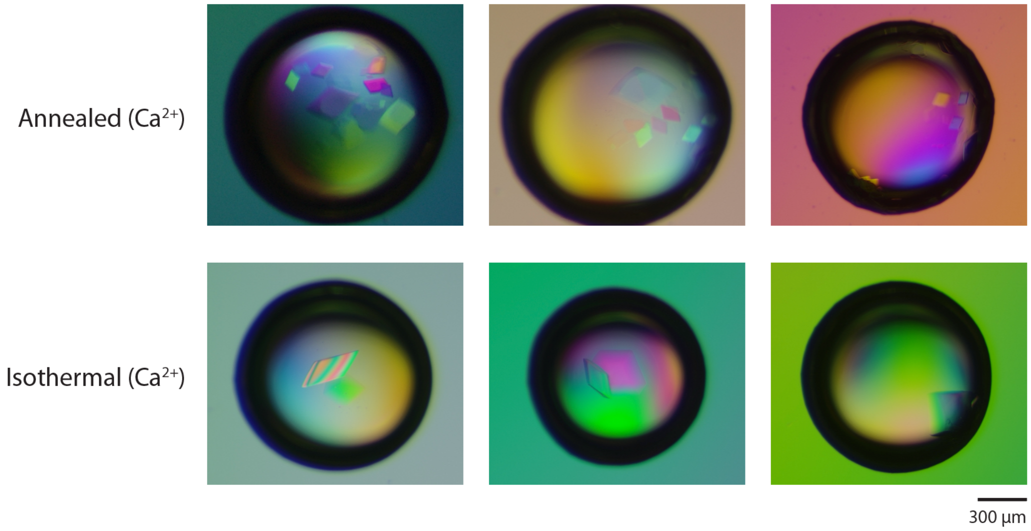


**Supplementary Figure 16**. Additional representative images of crystals grown from 3-turn tensegrity triangle motif annealed or isothermally assembled at 50 °C.

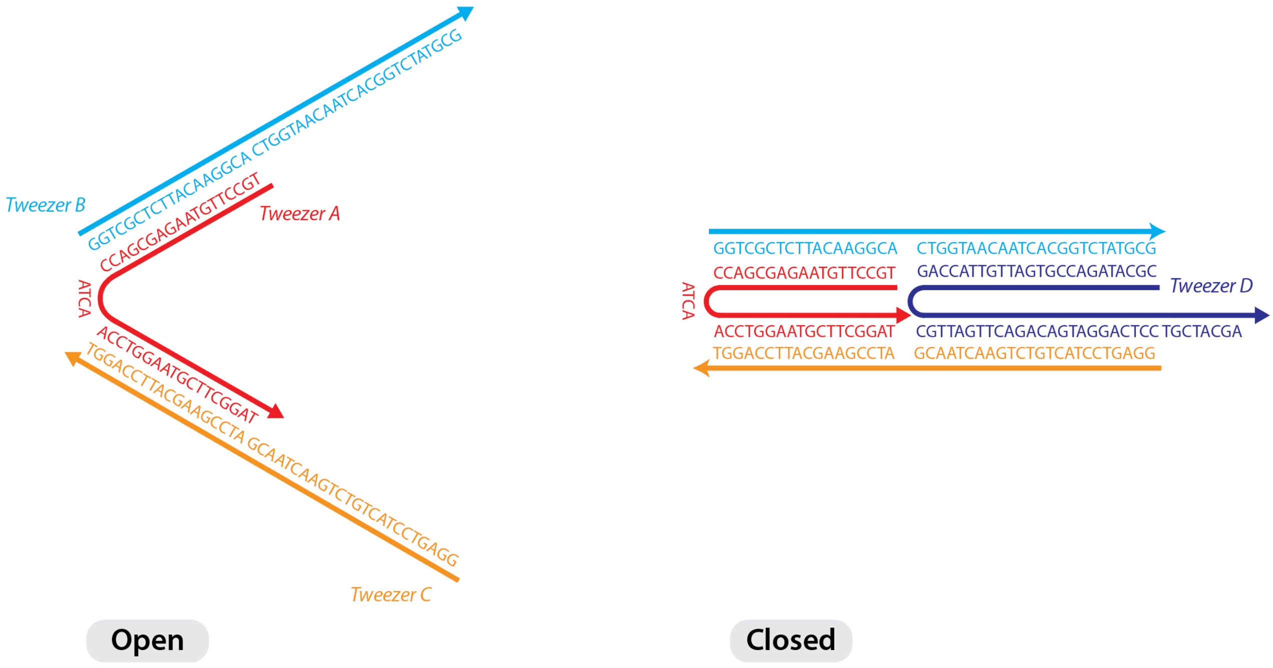


**Supplementary Figure 17**. Design and sequence of the DNA tweezer.

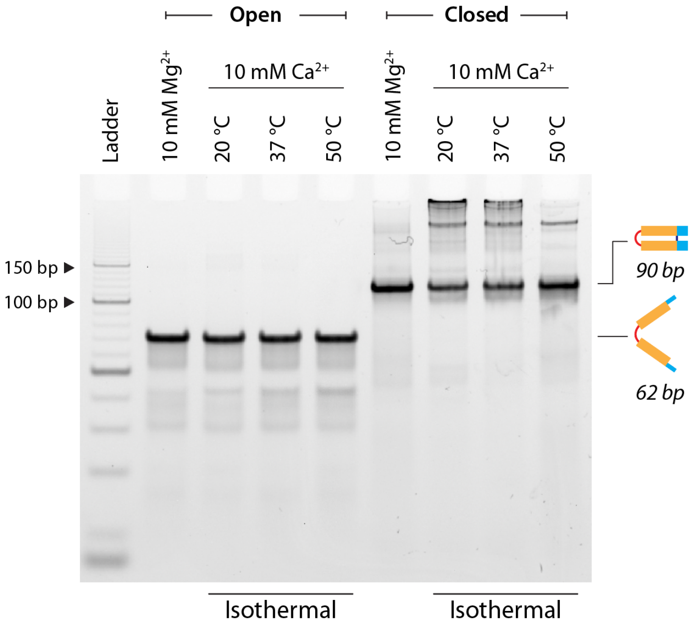


**Supplementary Figure 18**. Assembly of DNA tweezer in open and closed forms under isothermal conditions in 1× TAE buffer containing 10 mM $Ca^{2+}$. Lane 2 and 6 shows the control DNA tweezer, open and closed form respectively, assembled in $Mg^{2+}$ containing buffer using the annealing protocol.

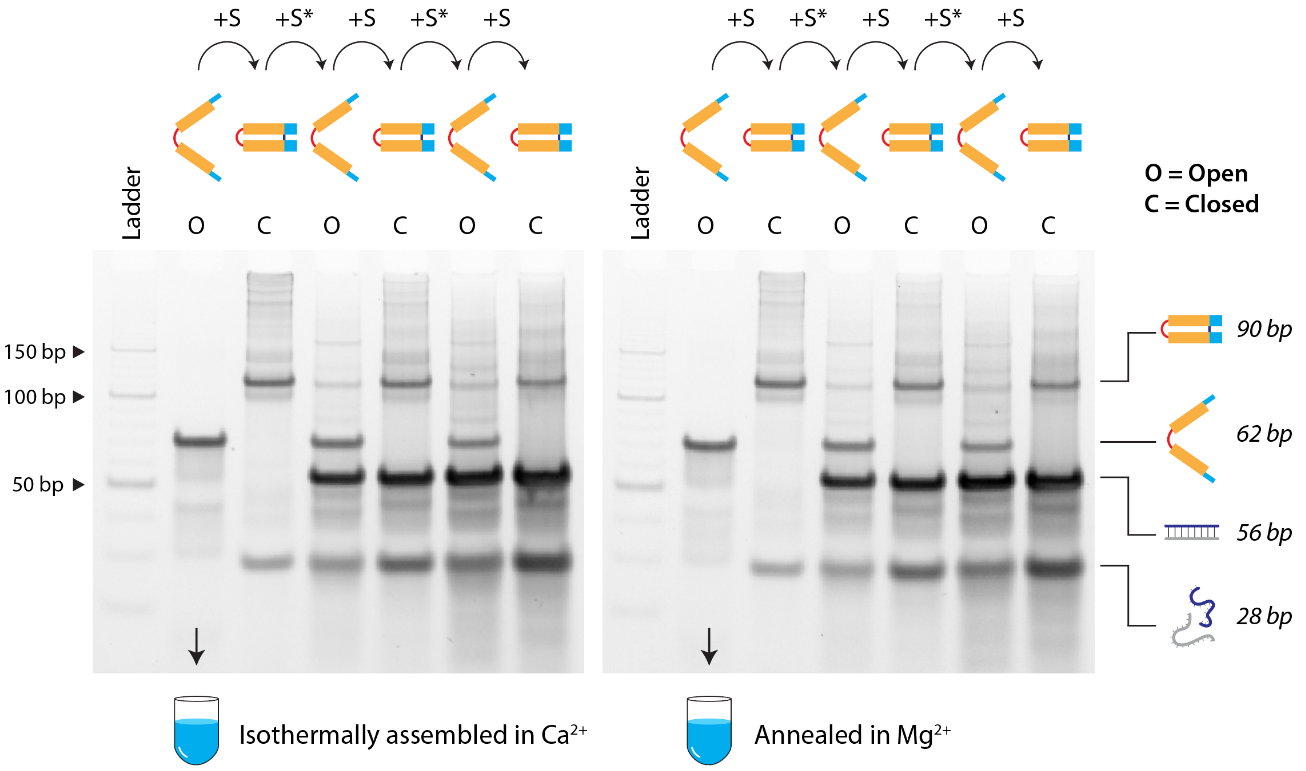


**Supplementary Figure 19**. Non-denaturing polyacrylamide gels showing repeated reconfiguration of the DNA tweezer between the open and closed states. Left: DNA tweezer was isothermally assembled in 1× TAE buffer containing 10 mM $Ca^{2+}$. Right: The tweezer was annealed 1× TAE-$Mg^{2+}$. Full gels of the images shown in Fig 5b.

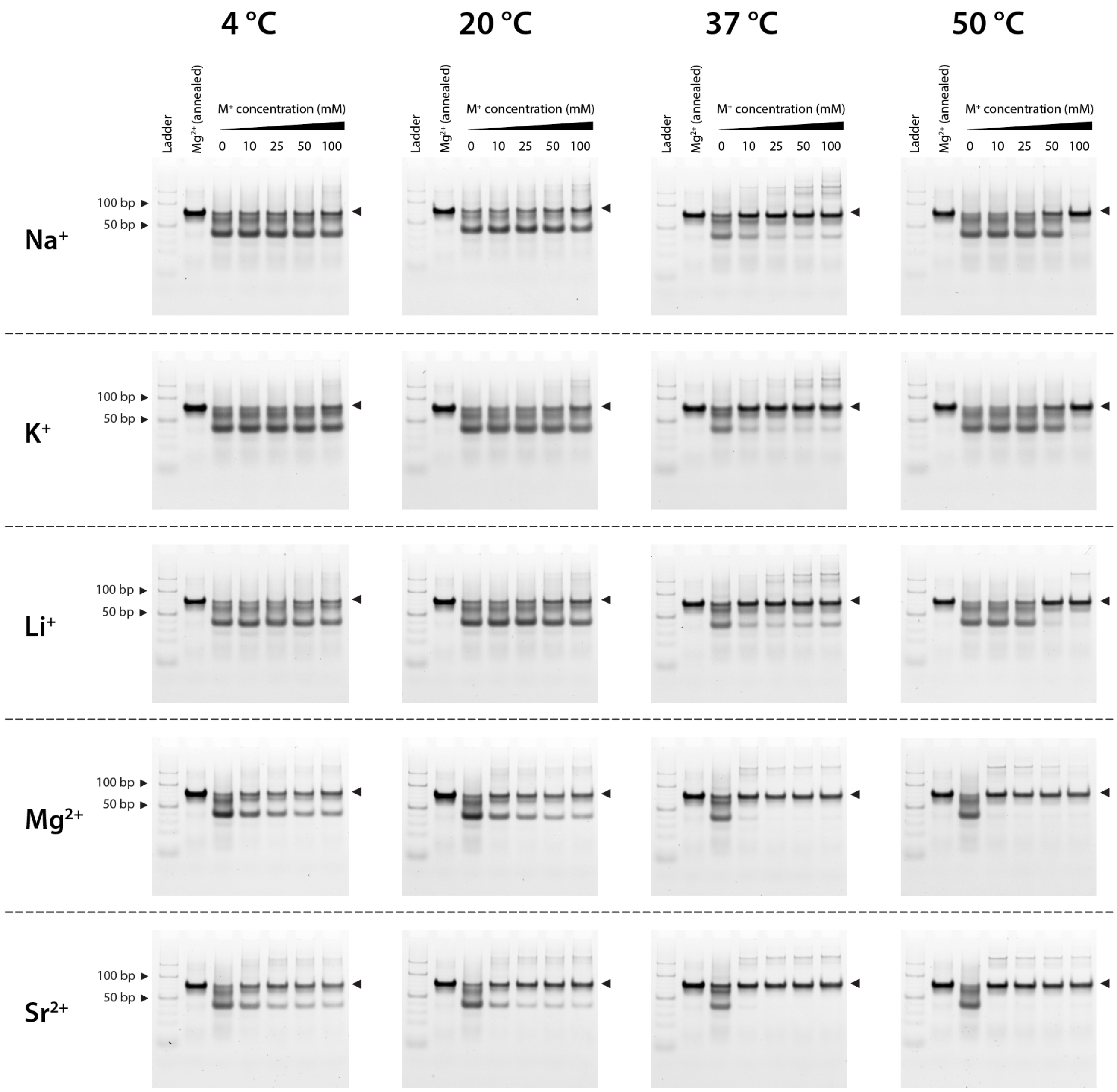


**Supplementary Figure 20.** Non-denaturing polyacrylamide gels showing isothermal assembly of DX-O in 1× TAE buffer containing different metal ions at 4 °C, 20 °C, 37 °C, and 50 °C. Lane 2 in all the gels shows DX-O assembled in $Mg^{2+}$ using the annealing protocol. Full gels of the images shown in Fig 6a-e.

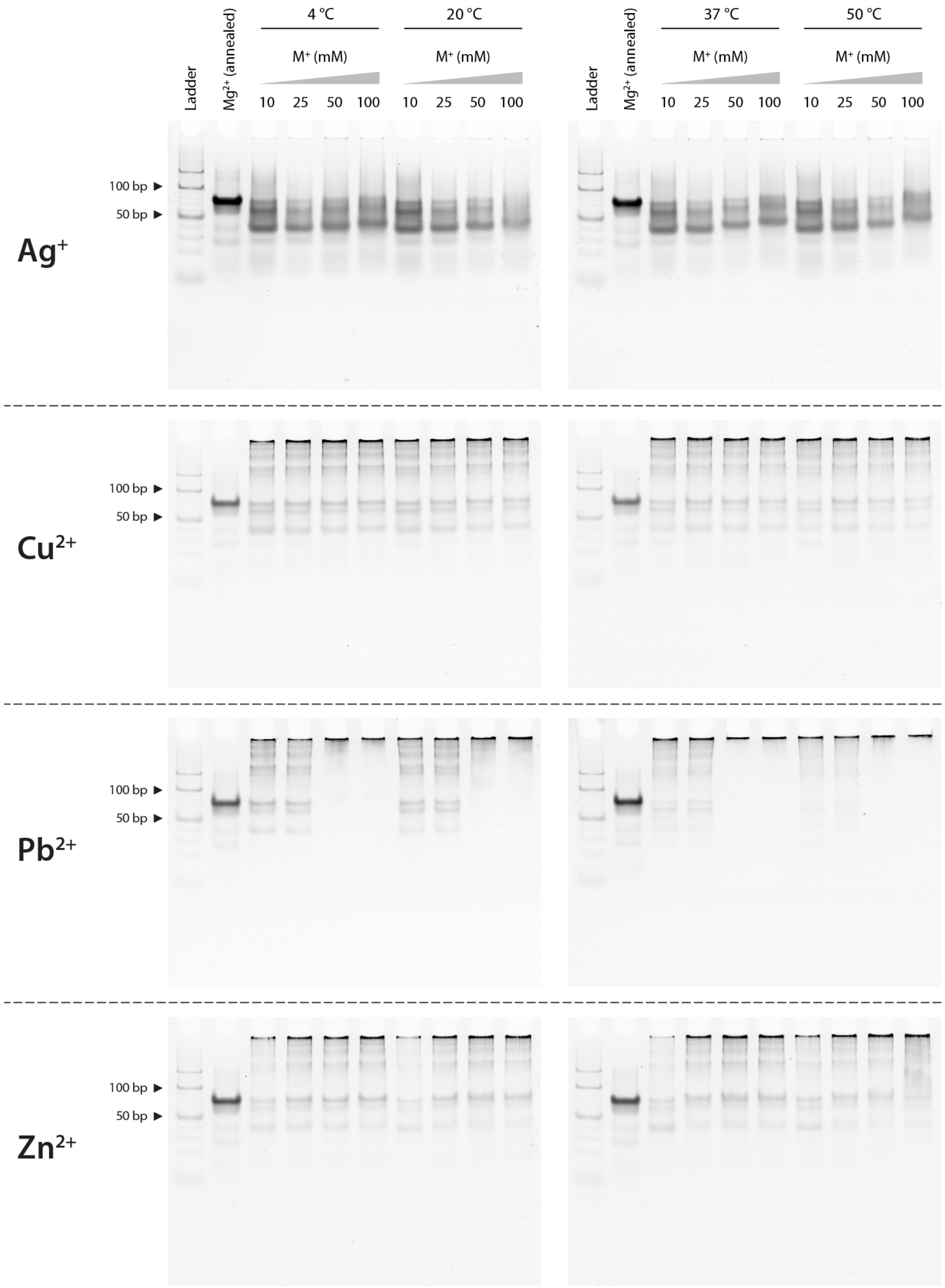


**Supplementary Figure 21**. Non-denaturing polyacrylamide gels showing poor isothermal assembly of DX-O in 1× TAE buffer containing different metal ions at 4 °C, 20 °C, 37 °C, and 50 °C. Lane 2 in all the gels shows DX-O assembled in $Mg^{2+}$ using the annealing protocol.

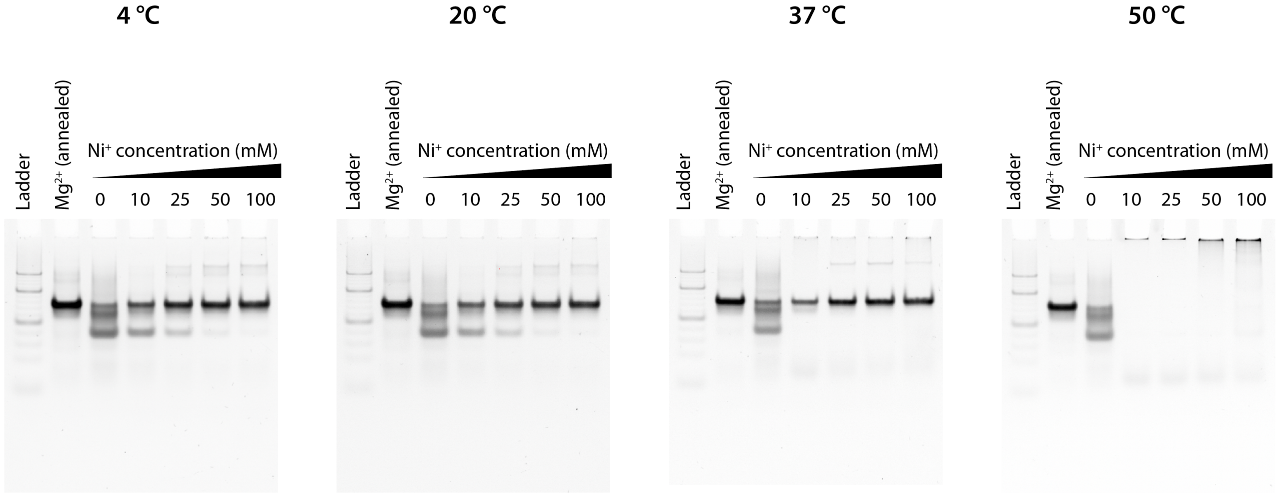


**Supplementary Figure 22**. Non-denaturing polyacrylamide gels showing the isothermal assembly of DX-O in 1× TAE buffer containing different concentrations of $Ni^{2+}$ at 4 °C, 20 °C, 37 °C, and 50 °C. Lane 2 in all the gels shows DX-O assembled in $Mg^{2+}$ using the annealing protocol. Full gels of images shown in Fig 6f.

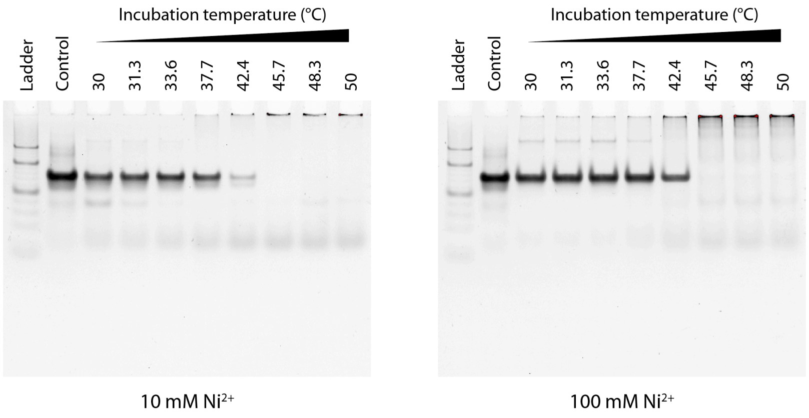


**Supplementary Figure 23**. Non-denaturing polyacrylamide gels showing effect of isothermal assembly temperature on the structural integrity of DX-O when assembled 1× TAE buffer containing 10 mM and 100 mM $Ni^{2+}$. Full gels of images shown in Fig 6g.

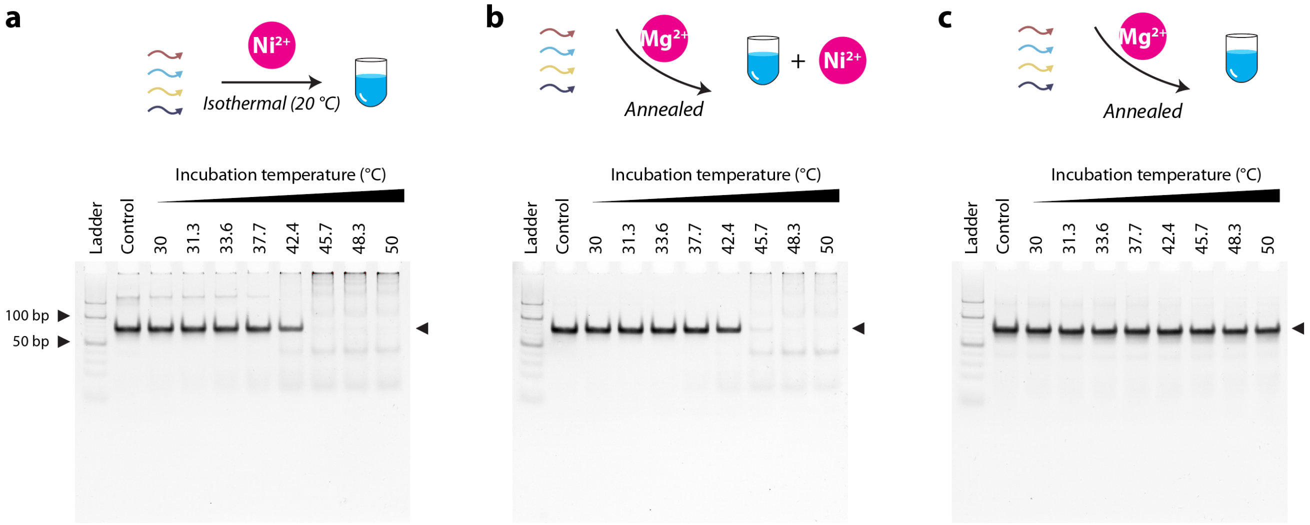


**Supplementary Figure 24**. Non-denaturing polyacrylamide gels showing the effect of temperature on the structural integrity of DX-O in 1× TAE buffer containing 100 mM $Ni^{2+}$. (a) DX-O was isothermally assembled in 1× TAE-$Ni^{2+}$ at 20 °C and incubated at different temperatures for 2 h. (b) DX-O was thermally annealed in 1× TAE-$Mg^{2+}$ and incubated with 100 mM $Ni^{2+}$ at different temperatures for 2 h. (c) DX-O was thermally annealed in 1× TAE-$Mg^{2+}$ and the solution was incubated at different temperatures for 2 h. Full gels of images shown in Fig 6h.

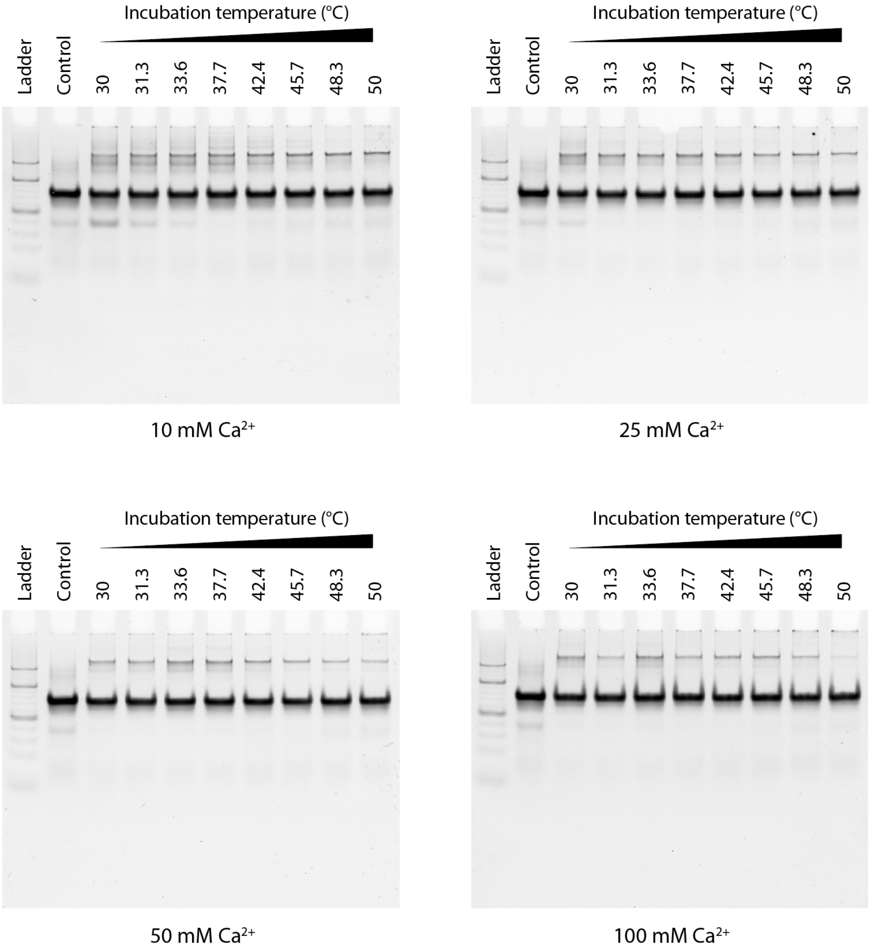


**Supplementary Figure 25**. Non-denaturing polyacrylamide gels showing the isothermal assembly of DX-O at different temperatures in 1× TAE buffer containing 10 mM, 25 mM, 50 mM, and 100 mM $Ca^{2+}$.

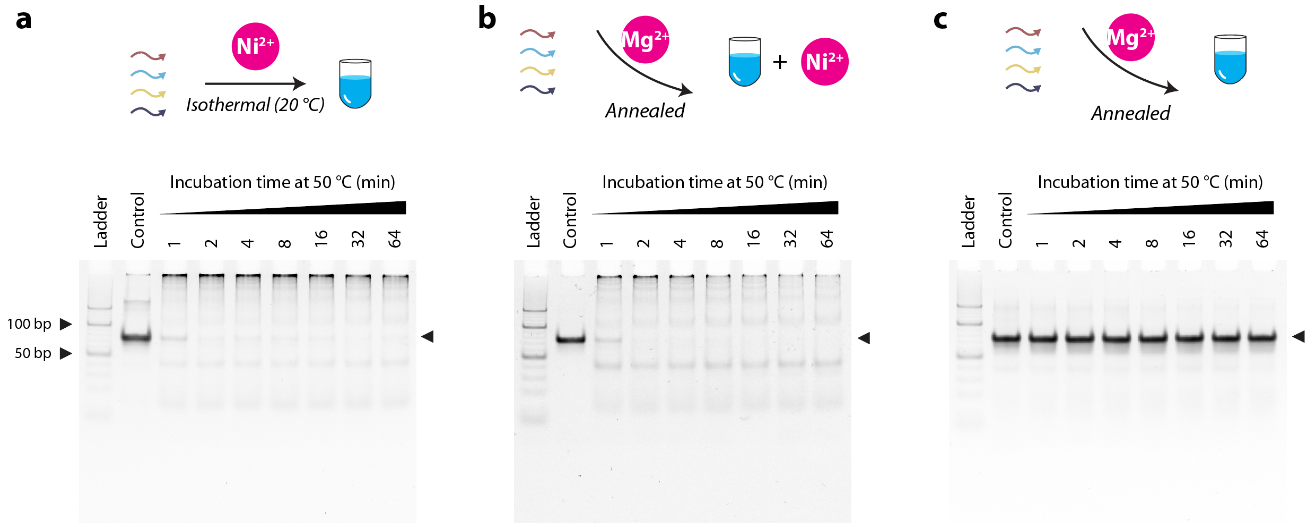


**Supplementary Figure 26**. Non-denaturing polyacrylamide gels showing the loss of structural integrity of DX-O in 1× TAE buffer containing 100 mM $Ni^{2+}$ over 64 min when (a) DX-O was isothermally assembled in 1× TAE-$Ni^{2+}$ at 20 °C, (b) DX-O was annealed in 1× TAE-$Mg^{2+}$ first and then mixed with $Ni^{2+}$, (c) DX-O was annealed in 1× TAE-$Mg^{2+}$ and incubated in the same buffer without any $Ni^{2+}$.

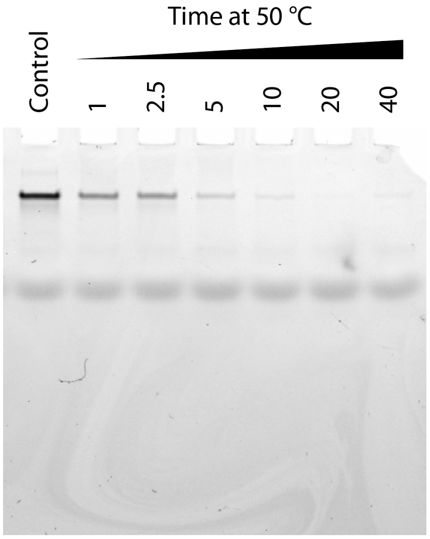


**Supplementary Figure 27**. Non-denaturing polyacrylamide gels showing degradation of single-stranded DNA when incubated in 1× TAE buffer containing 100 mM $Ni^{2+}$ for different time points at 50 °C.

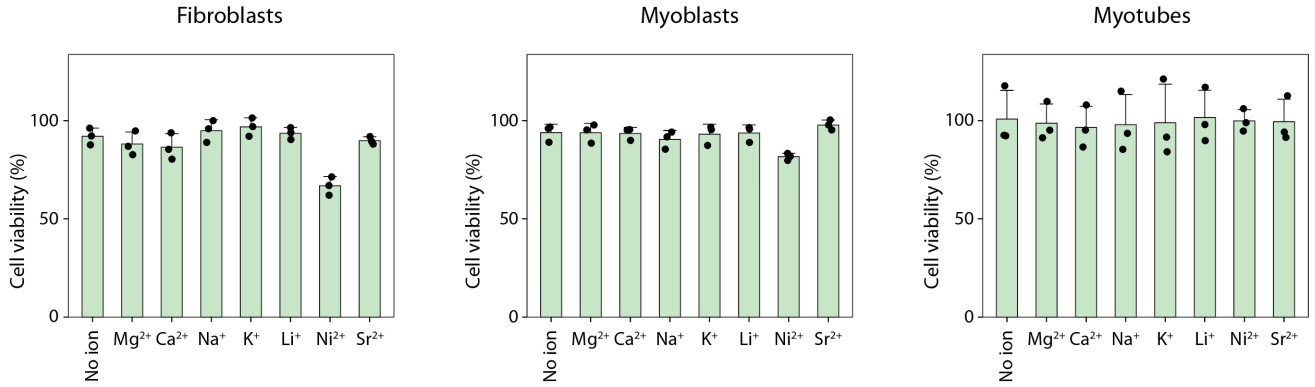


**Supplementary Figure 28**. Viability of fibroblasts, myoblasts and myotubes treated with 1× TAE containing different ions. Data is normalized to viability of untreated cells.

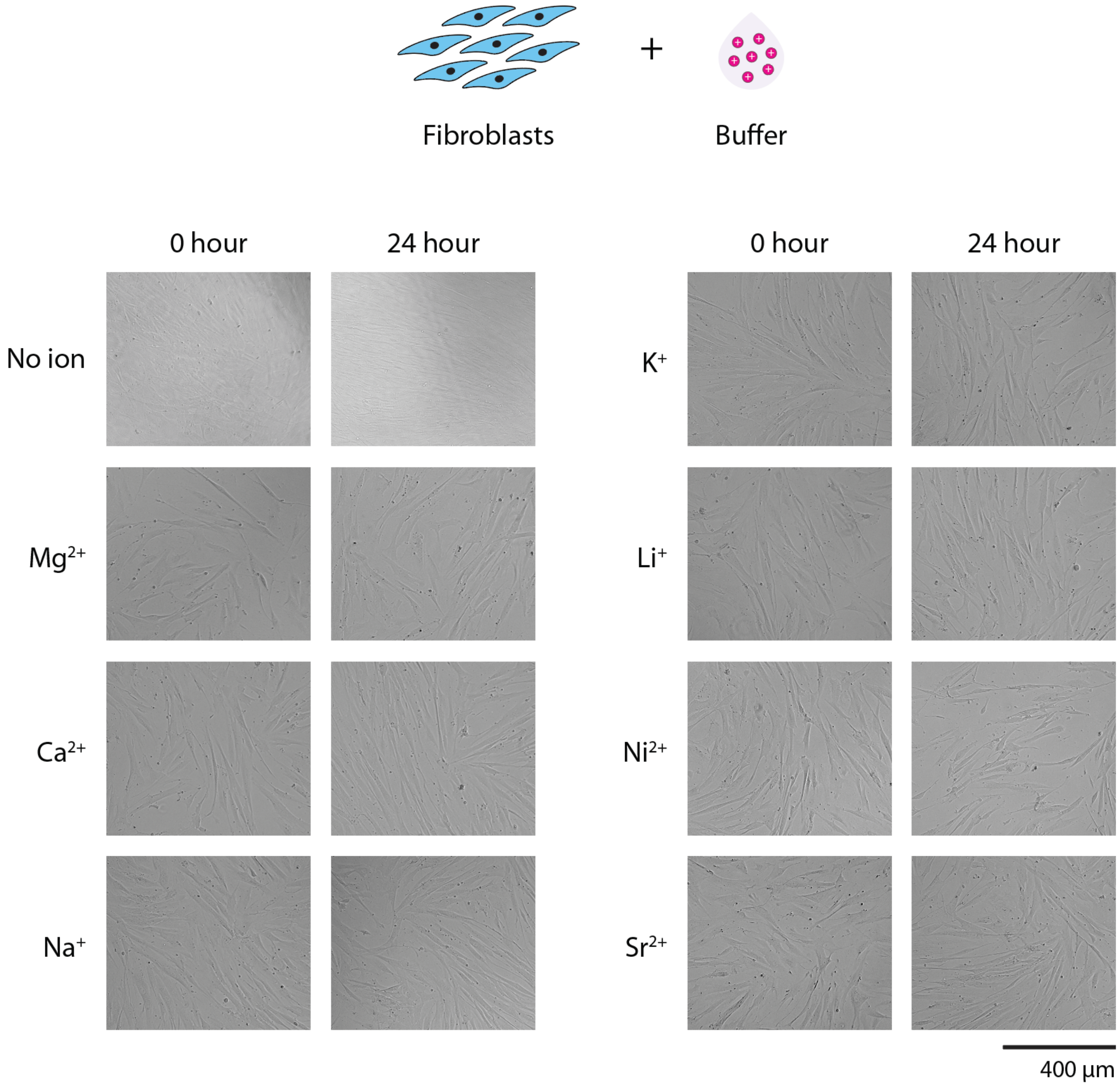


**Supplementary Figure 29**. Brightfield microscopy images of fibroblasts treated with 1× TAE buffer containing different ions.

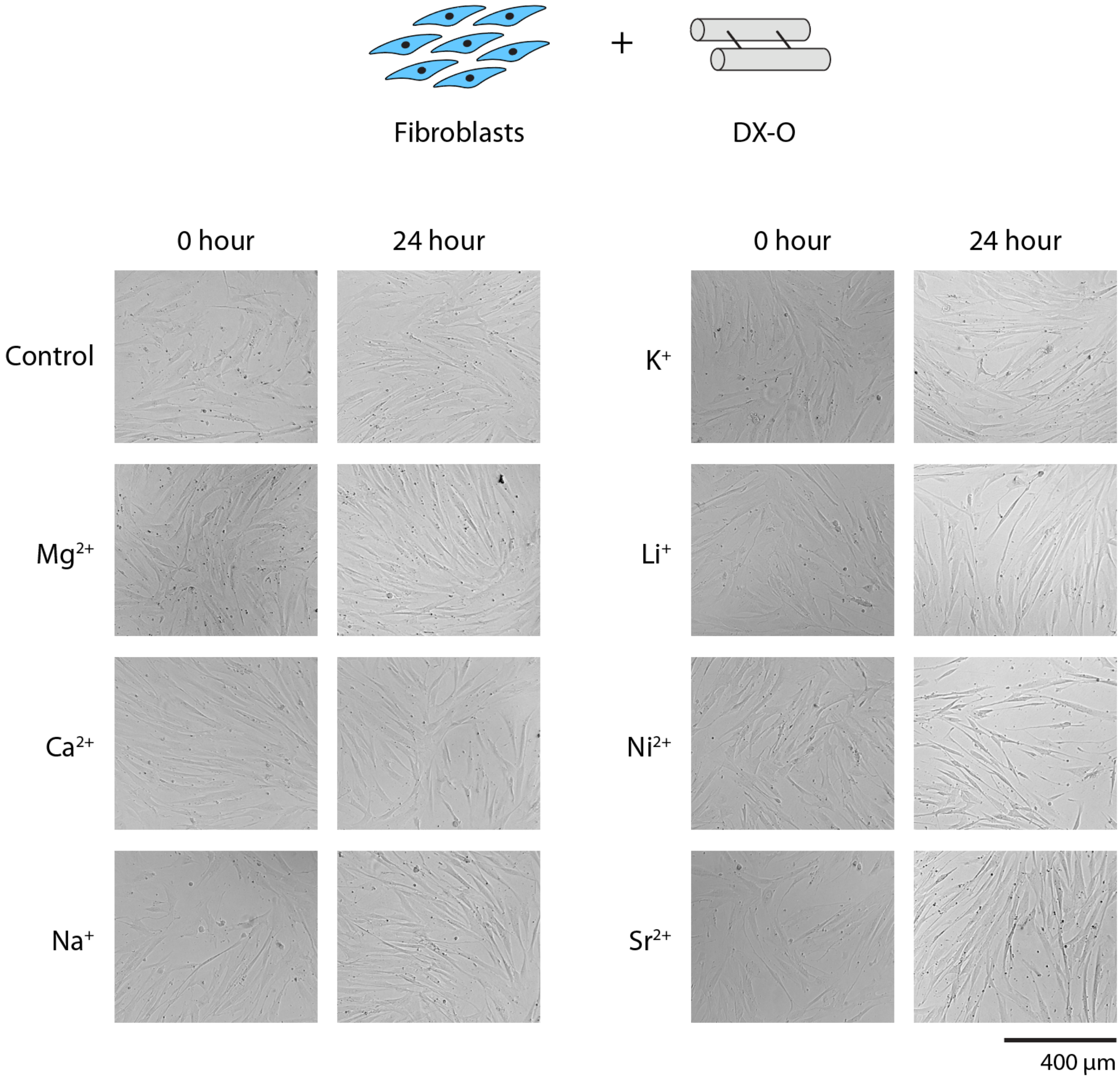


**Supplementary Figure 30**. Brightfield microscopy images of fibroblasts treated with DX-O motif isothermally assembled at 50 °C in 1× TAE buffer containing different ions. Control is DX-O motif assembled in $Mg^{2+}$ using thermal annealing.

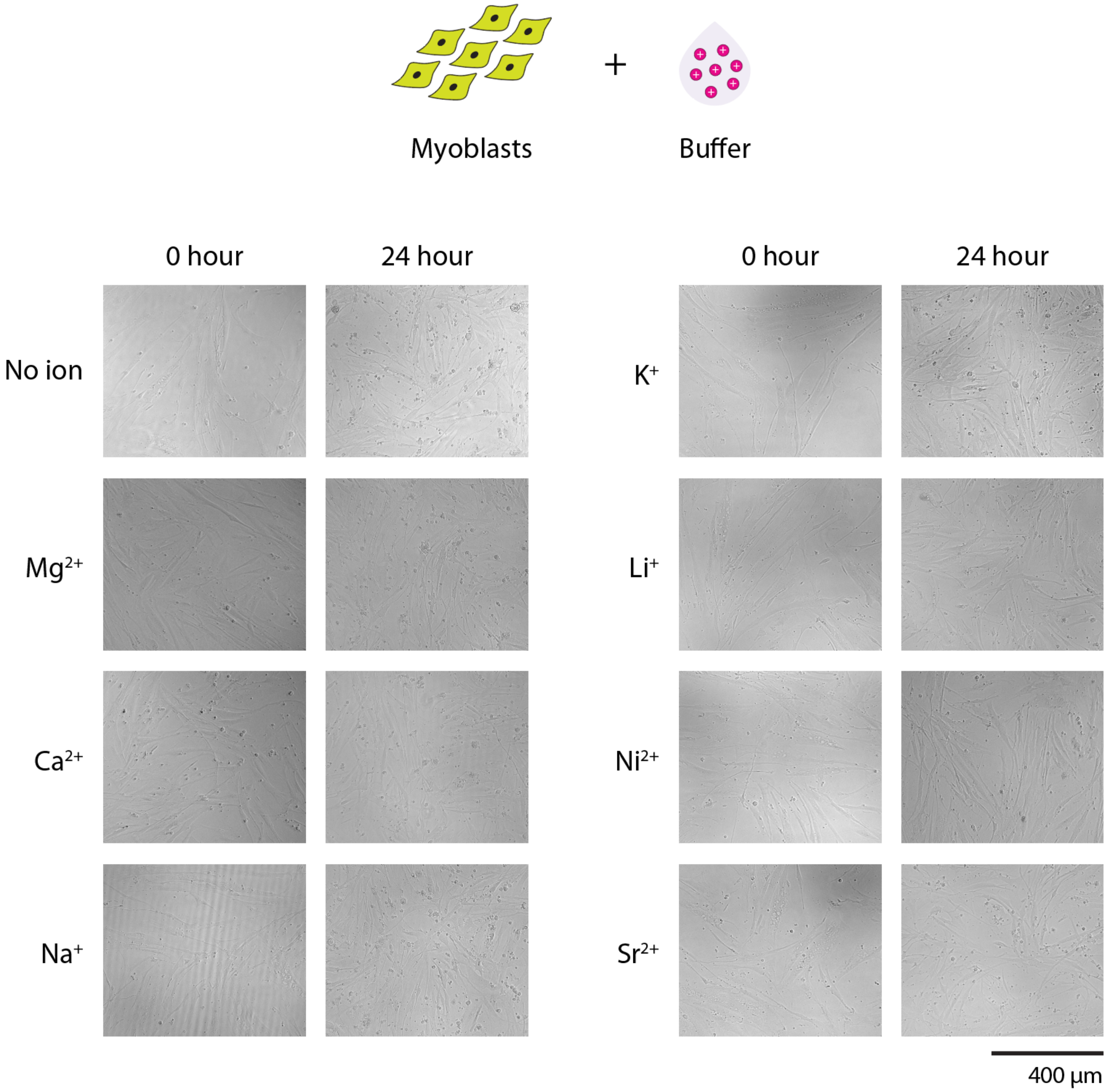


**Supplementary Figure 31**. Brightfield microscopy images of myoblasts treated with 1× TAE buffer containing different ions.

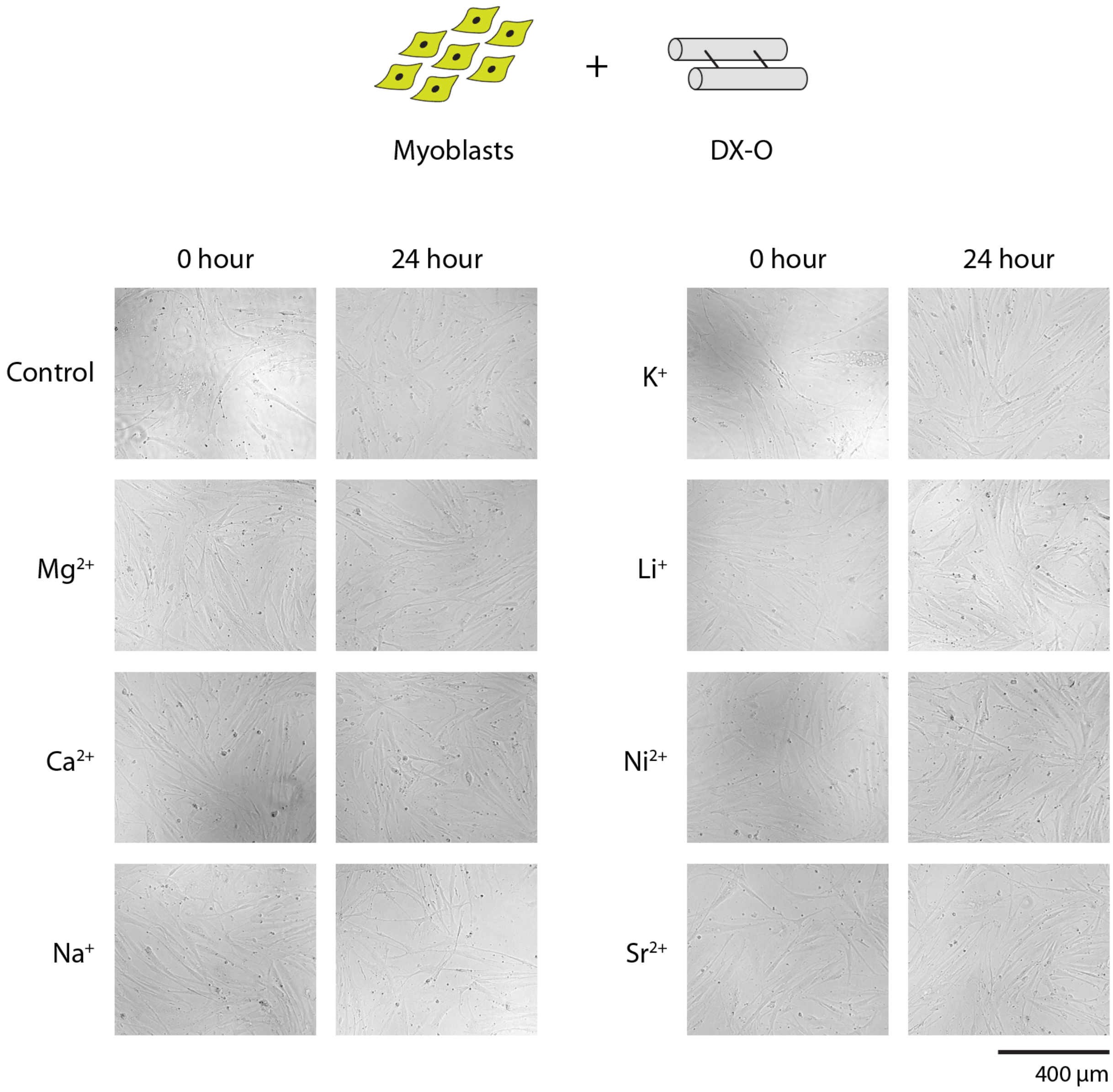


**Supplementary Figure 32**. Brightfield microscopy images of myoblasts treated with DX-O motif isothermally assembled at 50 °C in 1× TAE buffer containing different ions. Control is DX-O motif assembled in $Mg^{2+}$ using thermal annealing.

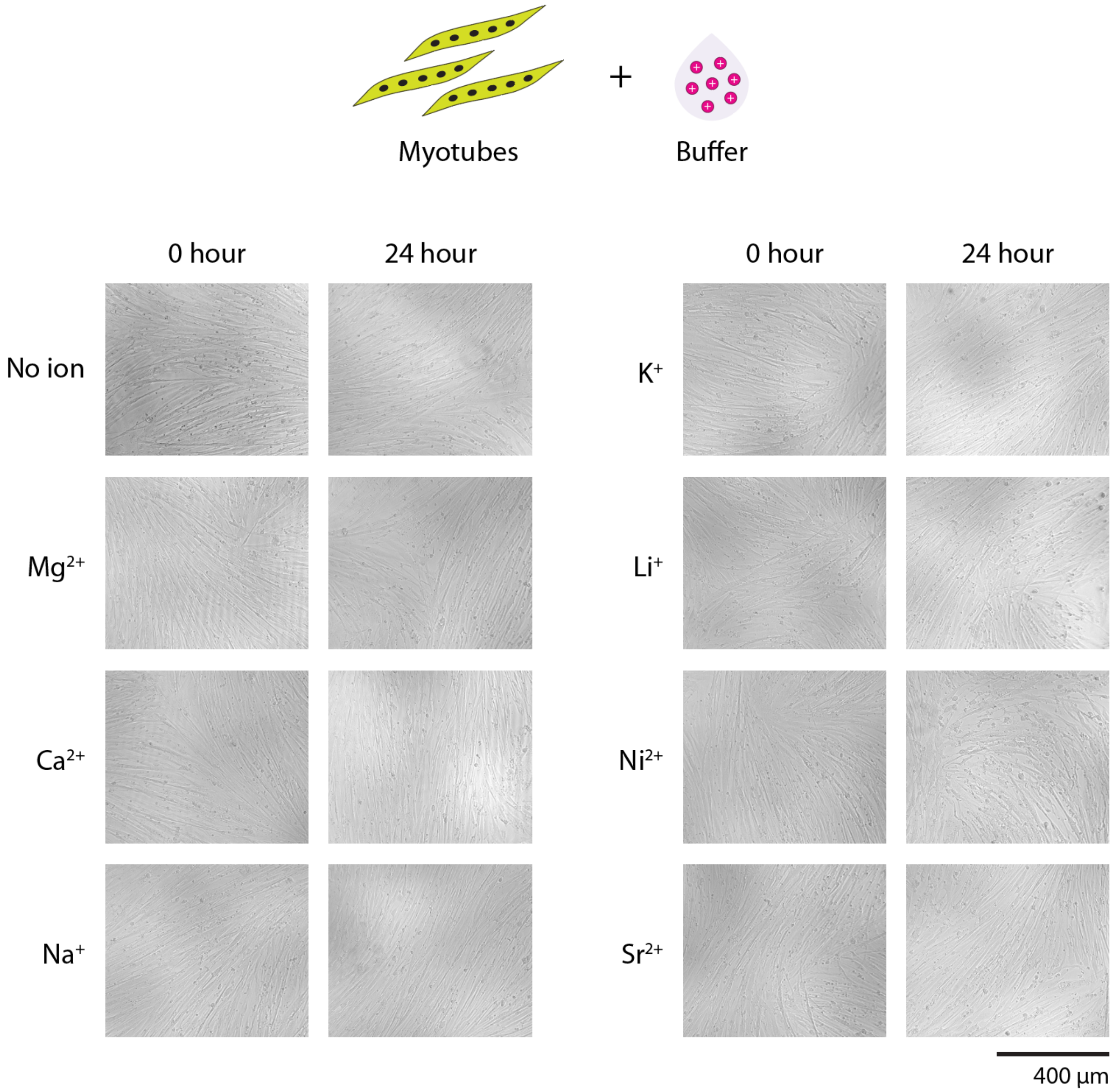


**Supplementary Figure 33**. Brightfield microscopy images of myotubes treated with 1× TAE buffer containing different ions.

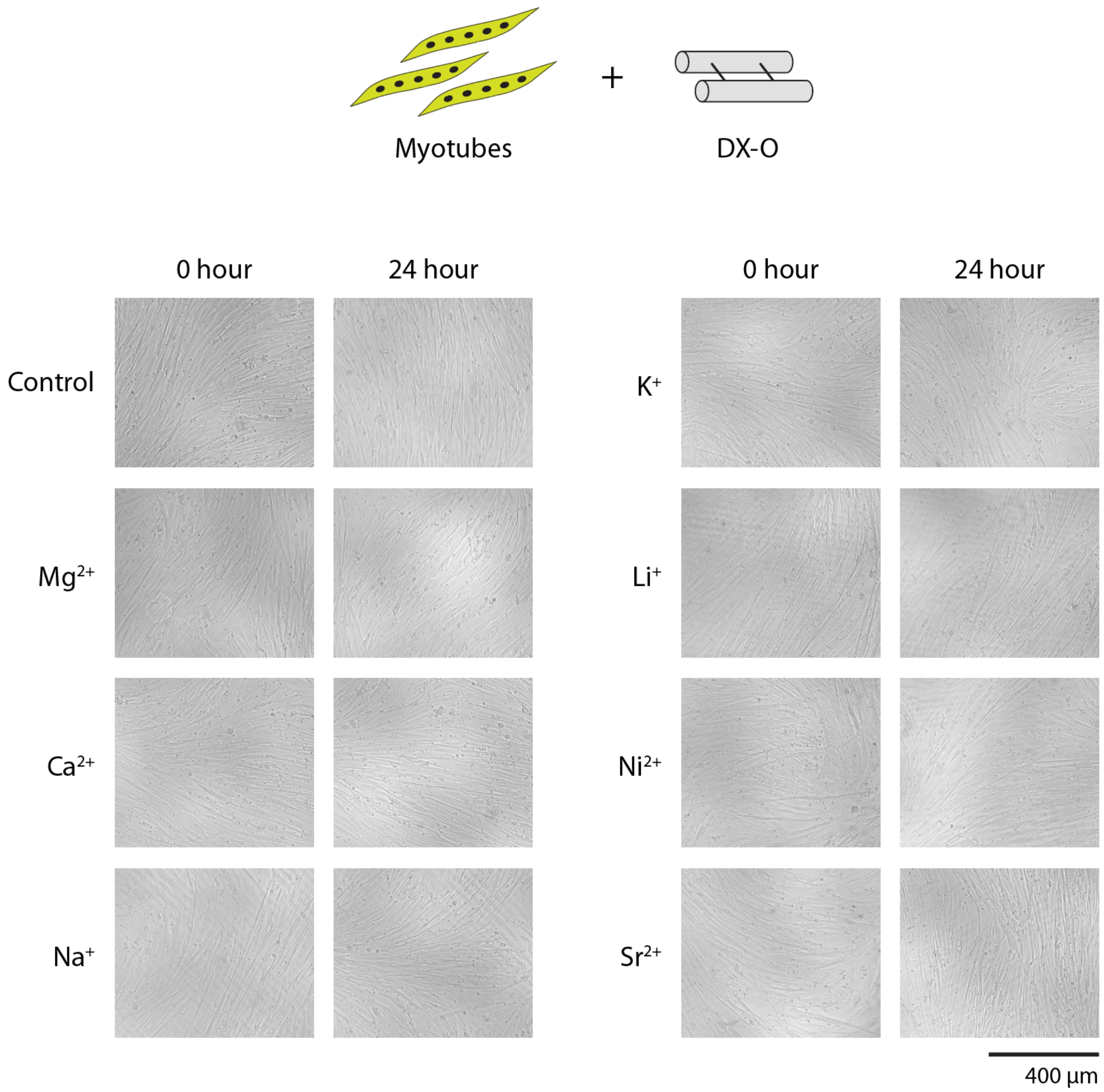


**Supplementary Figure 34**. Brightfield microscopy images of myotubes treated with DX-O motif isothermally assembled at 50 °C in 1× TAE buffer containing different ions. Control is DX-O motif assembled in $Mg^{2+}$ using thermal annealing.

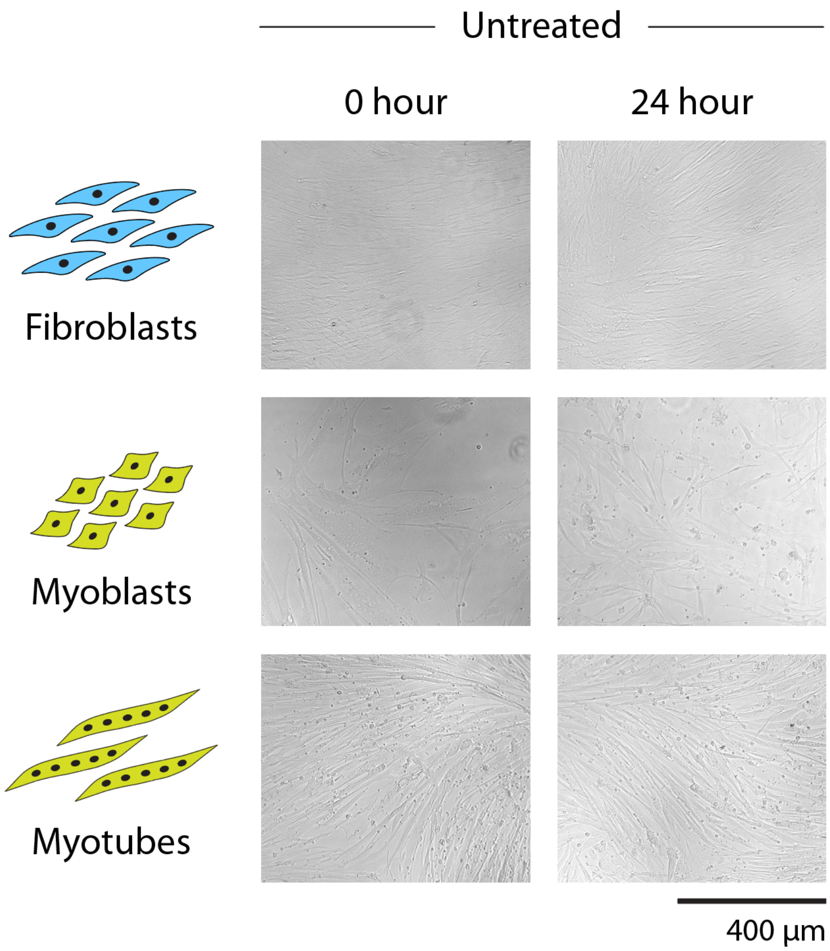


**Supplementary Figure 35**. Brightfield microscopy images of untreated control cells.

**Supplementary Table 1**. DX-O sequences (all sequences written from 5′ to 3′).

| | |
|---|---|
| DX-O-1 | CTCCATTCTGGACGCCATAAGATAGCACCTCGACTCATTTGCCTGCGGTAGAGT |
| DX-O-2 | AGACAGTAGCCTGCTATCTTATGGCGTGGCAAATGAGTCGAGGACGGATCGGTA |
| DX-O-3 | ACTCTACCGCACCAGAATGGAG |
| DX-O-4 | TACCGATCCGTGGCTACTGTCT |

**Supplementary Table 2**. 4-arm junction sequences.

| | |
|---|---|
| 4WJ-1 | CGCAATCCTGAGCACG |
| 4WJ-2 | CGTGCTCACCGAATGC |
| 4WJ-3 | GCATTCGGACTATGGC |
| 4WJ-4 | GCCATAGTGGATTGCG |

**Supplementary Table 3**. DX-E sequences.

| | |
|---|---|
| DX-E-1 | GCGACATCCTGCCGCTATGATTACACAGCCTGAGCATT |
| DX-E-2 | TTGATCGGACGATACTACATGCCAGTTGGACTAACGGC |
| DX-E-3 | GCCGTTAGTGGATGTCGC |
| DX-E-4 | AATGCTCACCGATCAA |
| DX-E-5 | TGTAGTATCGTGGCTGTGTAATCATAGCGGCACCAACTGGCA |

**Supplementary Table 4**. 3-helix motif sequences.

| | |
|---|---|
| 3HM-1 | ATCGAGAGACATAACTGCTTGACCACGCTGTATCGGAACCTGACTCCT |
| 3HM-2 | AGGAGTCACTCTCGAT |
| 3HM-3 | GGTATAGTATGCAACGTGAATGAACAAGGTGAGGTGTCAATGGAGAT |
| 3HM-4 | ATCTCCATTGACAGGTCAAGCAGTTATGTGGTTCTGCATACTATACC |
| 3HM-5 | ATAGCACCACTGCAAG |
| 3HM-6 | CTTGCAGTCCTTGTTCATTCACGTCGATACAGCGTCCTCAGGTGCTAT |

**Supplementary Table 5**. 4-helix motif sequences.

| | |
|---|---|
| 4HM-1 | CCACGTCTTA |
| 4HM-2 | AGTGTGAAACTCAGCAGATCCTATGCGTTC |
| 4HM-3 | GAACGCATAGTCGGCTGGGAAAATGCTTAGAAGATTAAGTTTGATCTCGTGG |
| 4HM-4 | TAAGAGCTGAATCATGGATTACTCGGATACGTTAAAGGTAGGGTTTCACACT |
| 4HM-5 | CTAGTAGTCCTCCGAGTAATCCATGATCCTACCTTTAACGTATTTGGTAATGGT |
| 4HM-6 | GGCTACAGACTGCATTTTCCCAGCCGAAAACTTAATCTTCTAAAATTCTCAAAC |
| 4HM-7 | ACCATTACCAAAGGACGAATTAGTCTGTAGCC |
| 4HM-8 | GTTTGATACTAG |

**Supplementary Table 6**. 3-turn tensegrity triangle motif sequences.

| | |
|---|---|
| 3TT-1 | GAGCAGCCTGTACTCGGCTTGTCGGACATCA |
| 3TT-2 | TCTGATGTGGCTGC |
| 3TT-3 | CCGAGTACACCGACAAGCCGAGTACACCGACAAGCCGAGTACACCGACAAG |

**Supplementary Table 7**. DNA tweezer sequences.

| | |
|---|---|
| Tweezer A | TGCCTTGTAAGAGCGACCATCAACCTGGAATGCTTCGGAT |
| Tweezer B | GGTCGCTCTTACAAGGCACTGGTAACAATCACGGTCTATGCG |
| Tweezer C | GGAGTCCTACTGTCTGAACTAACGATCCGAAGCATTCCAGGT |
| Tweezer D | CGCATAGACCGTGATTGTTACCAGCGTTAGTTCAGACAGTAGGACTCCTGCTACGA |
| Tweezer D comp | TCGTAGCAGGAGTCCTACTGTCTGAACTAACGCTGGTAACAATCACGGTCTATGCG |

**Supplementary Table 8**. The percentage of nucleotides with Root Mean Square Fluctuation (RMSF) values > 3 Å for each strand.

| Strand | $Mg^{2+}$ | $Ca^{2+}$ | $K^{+}$ | $Na^{+}$ |
|---|---|---|---|---|
| A | 62.3 | 11.3 | 100 | 100 |
| B | 39.6 | 15.1 | 73.6 | 75.5 |
| C | 76.2 | 0 | 100 | 100 |
| D | 23.8 | 14.3 | 28.6 | 33.3 |
